# Network geometry shapes the transformation of task representations across human cortex

**DOI:** 10.1101/2025.03.14.643366

**Authors:** Lakshman N.C. Chakravarthula, Takuya Ito, Alexandros Tzalavras, Michael W. Cole

## Abstract

Flexible cognition requires the brain to generalize across tasks while preserving task-specific information. Previous work showed that task representations are compressed in association cortex and expanded in sensory and motor systems, but the mechanisms underlying these transformations are unclear. Here we test whether the geometry of intrinsic brain connectivity constrains how representations change between cortical regions. We analyzed fMRI acquired during 16 diverse cognitive tasks together with activity flow modelling, finding that connectivity dimensionality predicts representational changes across cortex. Low-dimensional connectivity compressed task representations and increased similarities across tasks, whereas high-dimensional connectivity expanded representations and supported conjunctive coding of task features. Activity flow modelling reproduced the observed compression to expansion pattern along the sensory-association-motor hierarchy and generated representational geometries that more closely matched targets than sources, consistent with transformation rather than transfer of information. These findings identify network geometry as a systems-level principle that shapes cortical representations and supports flexible cognition.

## Introduction

The human brain’s ability to perform diverse tasks efficiently is a hallmark of cognitive flexibility. A key question in the neuroscience of cognitive flexibility is understanding how distributed neural circuits are recruited and reconfigured to navigate diverse task states ^1^. Recent studies have shed light on how different brain regions flexibly represent information during the performance of diverse tasks, revealing structured patterns of neural activity that adaptively support diverse cognitive demands ^2–6^. Along the sensory-to-motor cortical axis, task representations are first compressed, then expanded ^6^. Concretely, different task conditions may produce distinct high-dimensional patterns in visual cortex reflecting their unique sensory features (colors, shapes, words). In association cortex, representations compress – conditions that share abstract task-relevant features (e.g., ’respond left’ or ’inhibit response’) become more similar despite differing sensory details. In motor cortex, dimensionality increases again as abstract features must be translated into specific motor implementations: a ’respond left’ command could mean left index finger, left middle finger, or other motor specifics, each requiring distinct neural patterns. This suggests that sensory information is initially compressed across task conditions to facilitate transferrable representations, before expanding into representations that specify task-specific actions. This compression-then-expansion pattern is thought to provide the representational prerequisites for cognitive flexibility – the capacity to perform diverse tasks while maintaining task-specific performance quality^7^. Specifically, flexible performance requires representations that balance two competing demands: generalizability (sharing learned components across tasks to enable transfer) and separability (maintaining distinct implementations to prevent interference). This balance – compression enabling generalization, expansion enabling separability – is what we refer to as the representational prerequisites for cognitive flexibility. Compression in some regions may enable generalization by emphasizing shared abstract features across tasks, while expansion in other regions supports separability by maintaining task-specific sensory details and motor implementations. However, the systems-level mechanisms that generate this functional architecture remain unclear.

Prior studies have shown that different regions prioritize encoding different aspects of task conditions: some studies have observed regions representing both task-relevant and irrelevant variables ^8^, while others observed coding schemes supporting selective collapse of task-irrelevant information ^4,9,10^ or maintaining abstract representations ^11–13^. Other studies point toward representations encoding conjunctive combinations of variables in a non-linear mixed-selective manner ^14–16^. Consequently, some brain regions exhibit low-dimensional representations, while others represent task conditions in high-dimensional spaces ^17^. These variations in representational geometry suggest differing computational functions across brain regions.^7,17^

A central challenge in systems neuroscience is understanding how the brain’s network architecture enables flexible cognition. While local computations within brain regions certainly contribute to representational geometry, the role of inter-regional connectivity architecture remains underexplored. Recent theoretical work predicts that connectivity structure should systematically constrain neural representations^18–20^, yet empirical validation in biological circuits, particularly during diverse cognitive tasks, has been limited. Prior studies have demonstrated that task-evoked activity can be predicted using network connectivity^21,22^, and that distributed connectivity patterns can even predict localized functional specialization like category selectivity^23^. Building on these findings, we test whether connectivity architecture provides a systematic organizing principle for representational transformations across the cortical hierarchy. Specifically, does the geometric structure of inter-regional connectivity – whether connections are low- or high-dimensional –systematically shape how task representations compress or expand between brain regions?

To test this hypothesis, we build upon the activity flow modeling framework, which employs empirical functional connections to construct generative models to predict task-evoked activity patterns ^24–26^. In particular, this modeling framework proposes that task-evoked activity in target brain regions can be explained by activity “flowing” (propagating) from source regions to the target region via source-target connectivity. Here, we extend this framework to predict the representational geometry of brain regions using diverse task sets.

Specifically, we hypothesized that the dimensionality of intrinsic (resting-state) connectivity patterns systematically shapes the dimensionality of task representations in target regions (Fig. 1A). These are fundamentally different measures: connectivity dimensionality reflects the geometric structure of how regions connect during a single brain state (rest), while representational dimensionality reflects how a region’s activity varies across diverse task conditions. Testing whether these independently-measured properties are systematically related would reveal whether network architecture constrains representational geometry. Critically, this prediction is non-trivial: low-dimensional connectivity could theoretically lead to representational expansion in target regions, if target regions are dominated by local, nonlinear processes, and high-dimensional connectivity could reflect random connection patterns bearing no systematic relationship to representations. Moreover, it is often assumed that inter-region connectivity plays a passive role (like inter-computer connectivity in the internet), wherein connections simply carry information, and any information transformation occurs within (rather than between) regions. Finding systematic relationships between connectivity geometry and representational transformations would demonstrate that network architecture provides structured constraints on neural computation.

**Figure 1:**
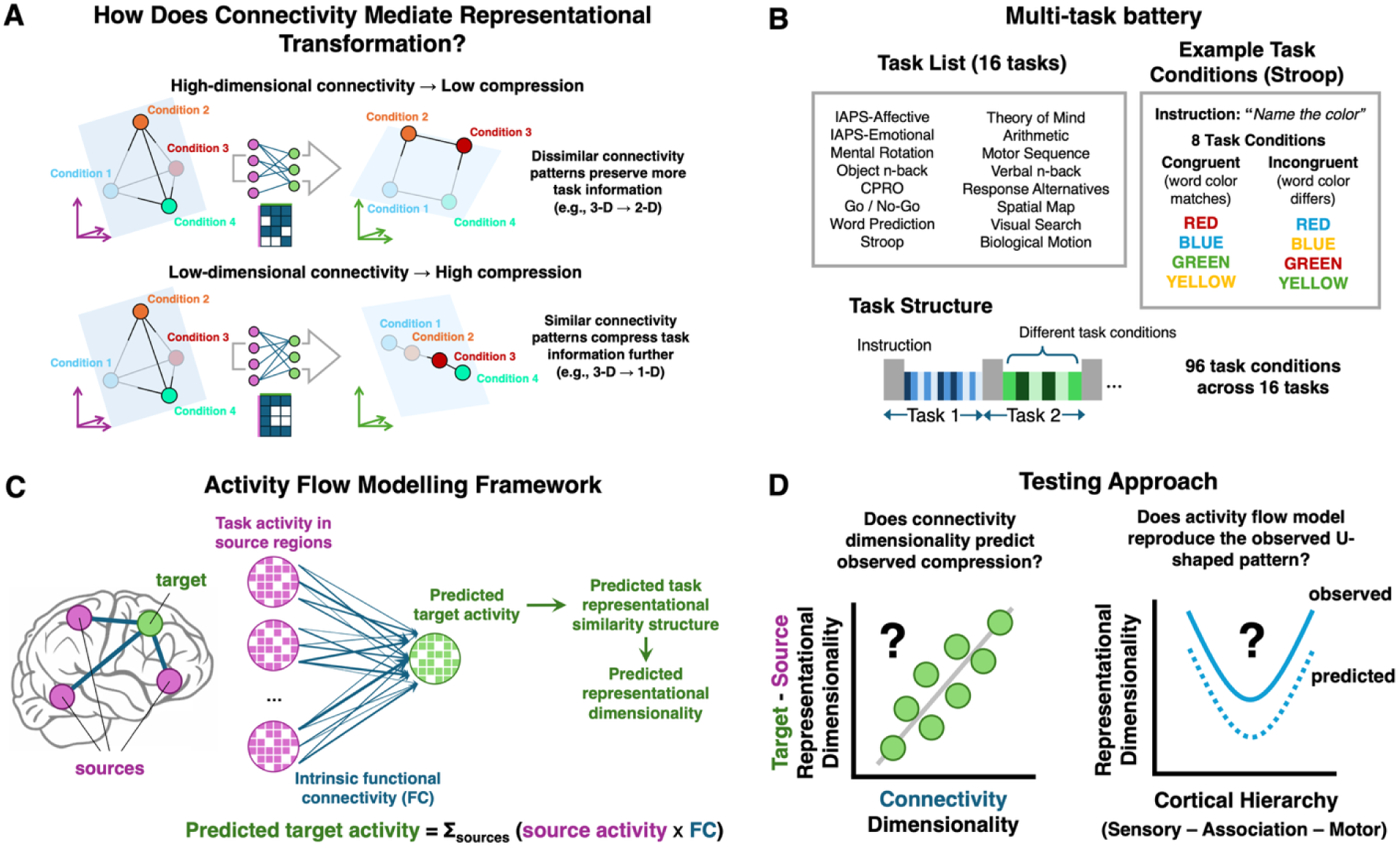
Testing how connectivity geometry shapes representational transformations across cortex. A) Hypothesis: Connectivity dimensionality determines compression versus expansion of task representations. Task conditions (colored spheres labeled 1-4) occupy positions in neural activity space. Representational dimensionality determines which pairwise task groupings can be separated by linear decision boundaries (transparent planes). Higher dimensionality enables decoding of more diverse task groupings: in 3D space, all three pairwise splits are linearly separable (1-2 vs 3-4; 2-3 vs 1-4; 2-4 vs 1-3); in 2D space, two pairwise splits remain possible (1-2 vs 3-4; 2-3 vs 1-4); in 1D space, only one pairwise split is possible (1-2 vs 3-4). *Top:* High-dimensional connectivity (dissimilar connectivity patterns among target vertices, shown in connectivity matrix) preserves representational dimensionality when source activity flows to target (3D→2D example), maintaining decodability of multiple task groupings. *Bottom:* Low-dimensional connectivity (similar patterns among target vertices) compresses representations (3D→1D example), eliminating the capacity to decode most task groupings. B) Multi-task battery provides diverse representational structure. *Left:* List of 16 active visual tasks spanning cognitive control, memory, language, and motor domains. *Right:* Example task structure using the Stroop task, which includes 8 task conditions crossing word-color congruency (congruent: word matches color; incongruent: word differs from color) with four ink colors. *Bottom:* Each task consisted of a 5-second instruction period followed by 30 seconds of task performance. Across all 16 tasks, we estimated activity patterns for 96 distinct task conditions using regularized GLM, providing rich representational diversity for measuring dimensionality. C) Activity flow modeling framework. Task activity in source regions (all functionally connected regions identified via sparse graphical lasso; see Methods) is projected to each target region via intrinsic (resting-state) functional connectivity weights estimated at the vertex level using principal component regression. For each task condition, predicted target activity is computed as the weighted sum of source activities: predicted target activity = Σ_sources_ (source activity × FC). From predicted target activity patterns across all 96 task conditions, we derive the predicted representational similarity structure (RSM) and predicted representational dimensionality using participation ratio of the eigenspectrum (see Methods). D) Testing approach. We test two key predictions. *Left:* Does connectivity dimensionality (measured from resting-state FC using participation ratio of the singular value spectrum; see Methods) predict the observed change in representational dimensionality from sources to target (compression: target dimensionality < average source dimensionality; expansion: target dimensionality > average source dimensionality)? Each point represents one cortical region. *Right:* Does the activity flow model reproduce the compression-then-expansion pattern of representational dimensionality observed along the cortical hierarchy from sensory through association to motor regions? Solid line shows observed pattern (averaged across regions within each system for each subject); dotted line shows predicted pattern from activity flow model. (End of Figure 1 caption)

Here we examine how connectivity dimensionality shapes inter-areal representational transformations during diverse task performance. Using fMRI data from participants performing 16 different tasks (Fig. 1B), we first demonstrate a robust compression-then-expansion pattern in representational dimensionality across the cortical hierarchy and validate its generalizability in an independent Human Connectome Project dataset (n=352). We find that the dimensionality of intrinsic connectivity systematically predicts how task representations are transformed between brain regions: low-dimensional connectivity leads to compression of representations, while high-dimensional connectivity leads to expansion. Through activity flow modeling, we show that these connectivity patterns actively generate observed representational geometries across the cortex. Moreover, we demonstrate through simulation that strong non-linear local computations can fully decouple connectivity dimensionality from representational transformations. Rather than implementing transformations through complex local computations in each region, the brain leverages network architecture itself as a computational substrate. Regions with lower-dimensional connectivity exhibit stronger cross-task similarities, suggesting shared latent features that enable generalization across tasks, while expansions from high-dimensional connectivity support conjunctive coding of task variables. Higher representational dimensionality, particularly in sensory and association regions, predicts individual differences in general fluid intelligence. Together, these results reveal how the network geometry of the brain’s intrinsic connectivity architecture shapes the transformation of task representations to support flexible cognition.

## Results

### Representational compression (expansion) is associated with low- (high) dimensional FC

Our main hypothesis was that intrinsic connectivity shapes cross-region representational transformations across diverse tasks. This hypothesis is based on the activity flow modeling framework, which proposes that a region’s task-evoked activity is shaped by activity flowing from connected source regions, weighted by their connectivity strengths^25^. Consistent with this, our hypothesis predicted that there should be a relationship between cross-region representational dimensionality (compression or expansion, where compression indicates lower-dimensional representations in the target relative to source regions, and expansion indicates higher-dimensional representations) and connectivity dimensionality. We expected to observe that low connectivity dimensionality between regions compressed task representations, and that high connectivity dimensionality expanded task representations.

We examined representational transformations between connected regions using a two-step approach to FC estimation (see Methods). In brief, we estimated a subject-specific FC graph from source vertices (all vertices of non-target regions connected to the target region) using graphical lasso (to identify connected region pairs) and principal components regression (to estimate vertex-level connectivity weights within those pairs). This whole-cortex vertex-level FC formed the basis for our subsequent analyses of connectivity dimensionality and its relationship to representational compression.

We measured representational dimensionality in each region by computing the participation ratio of the eigenspectrum of the cross-validated task representational similarity matrix (RSM). The participation ratio quantifies how many dimensions are needed to capture the variance in neural representations. Higher values indicate that task conditions produce distinct activity patterns across many dimensions, while lower values indicate that patterns can be described using fewer dimensions. Intuitively, if the 96 task conditions were represented as completely distinct patterns (orthogonal to each other), they would span a near-96-dimensional space (in a region with number of vertices >> 96). However, if some conditions share similar patterns (e.g., tasks requiring similar responses), they can be represented in a lower-dimensional subspace. As in prior work^6^, we observed that representational dimensionality strongly correlates with decodability of task conditions from one another (Pearson’s r = 0.755, spatial autocorrelation (SA)-controlled permutation test: p_perm_ < 0.001), thus reflecting the granularity of task information in the representations. We defined compression as the difference between the representational dimensionality of the target region and the average representational dimensionality of its source regions (positive values indicate expansion). This allowed us to quantify whether the target region’s representation was more compressed (lower dimensionality) or expanded (higher dimensionality) relative to its sources. For connectivity dimensionality, we sought a measure that would parallel our representational dimensionality metric while accounting for the unique structure of connectivity data between source and target regions. We therefore performed singular value decomposition on the connectivity matrix – an established approach for analyzing rectangular matrices – and computed the participation ratio of its singular value spectrum. As with representational dimensionality, this yields a continuous measure where higher values indicate more distinct components in the source-to-target connectivity structure.

Connectivity dimensionality reflects the similarity structure of source-to-target connectivity patterns rather than connection sparsity: it was strongly related to the mean pairwise correlation among target vertices’ connectivity patterns (r = −0.79, p_perm_ < 0.001) but unrelated to connection sparsity (p > 0.05; Extended Data Fig. 1).

Using these metrics, we tested whether changes in representational dimensionality from sources to each target correlated with the dimensionality of the intermediate FC. Building on prior work^6^, we plotted transformations along an FC gradient – a continuous axis derived from the principal structure of the whole-brain connectivity matrix, which runs from sensory through association to motor cortex and captures the cortical processing hierarchy (Fig. 2A), given the intuition that activity flow-based transformations likely occur along a sensory-association-motor axis during sensory-motor tasks. As expected, we observed a trend of compression-then-expansion in task representations along the FC gradient (Fig. 2B,D). As targets, sensory regions typically showed expansion, association regions exhibited compression, and motor regions expansion, forming a U-shaped trajectory of representational compression/expansion. This pattern can be understood through a concrete example: In visual cortex, different task conditions produce distinct high-dimensional patterns reflecting their unique sensory features (colors, shapes, words, spatial locations). In association regions, representations compress—conditions that share abstract task-relevant features (e.g., ’respond left’ or ’inhibit response’ or ’remember this stimulus’) become more similar despite differing sensory details. In motor cortex, dimensionality increases again because the abstract association-level representations must be translated into specific motor implementations: a ’respond left’ command could mean left index finger, left middle finger, left hand grip, or other motor specifics; a visual attention task and a working memory task might both require ’maintain fixation’ but with different timing parameters. Model comparisons confirmed the U-shape for representational dimensionality change (quadratic favored over linear, ΔAIC = 3.94, p_perm_ = 0.001).

**Figure 2:**
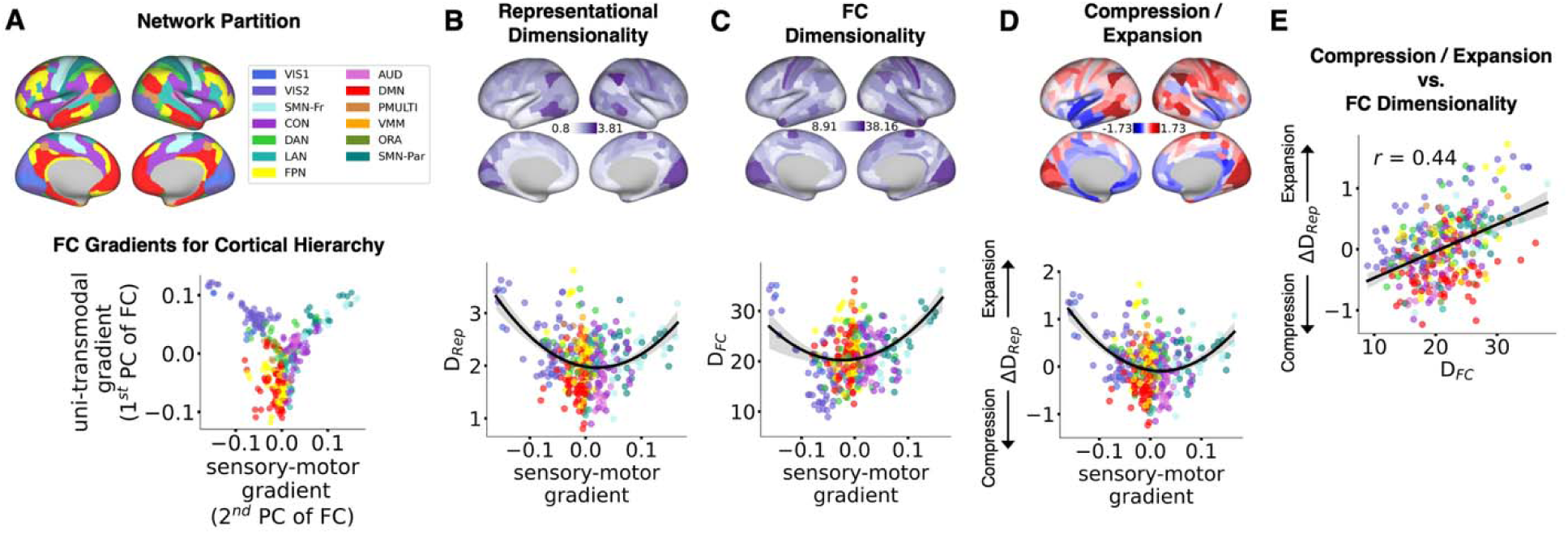
Connectivity dimensionality predicts representational compression and expansion across cortex. A) Cortical organization characterized by discrete networks and continuous gradients. *Top:* Cortical regions colored by functional network identity according to the Cole-Anticevic Brain Network Partition^30^, dividing cortex into 12 networks including visual (VIS1, VIS2), somatomotor (SMN-Fr: frontal/motor, SMN-Par: parietal/somatosensory), cingulo-opercular (CON), dorsal attention (DAN), language (LAN), frontoparietal (FPN), auditory (AUD), default-mode (DMN), posterior-multimodal (PMULTI), ventral-multimodal (VMULTI), and orbito-affective (ORA) networks. *Bottom:* Continuous FC gradients derived from principal components analysis of the group-averaged, thresholded parcel-level connectivity graph. Each dot represents one of 360 cortical regions, colored by network identity. The first principal component (y-axis) captures a uni-transmodal gradient separating sensory/motor regions from association regions. The second principal component (x-axis) captures a sensory-motor gradient running from sensory through association to motor regions, providing a continuous measure of cortical hierarchy position. We use this sensory-motor gradient (2nd PC) for visualization throughout. B) Representational dimensionality shows compression-then-expansion along the cortical hierarchy. *Top:* Brain maps showing representational dimensionality (DRep, participation ratio of task RSM eigenspectrum) for each region, averaged across subjects. *Bottom:* Representational dimensionality plotted along the sensory-motor gradient, with regions colored by network. Sensory regions (left) and motor regions (right) show high dimensionality, while association regions (center) show compressed representations, forming a U-shaped pattern (black curve: quadratic regression fit with 95% bootstrap CI as grey error band, 1000 iterations; quadratic model favored over linear, ΔAIC = 3.94, quadratic coefficient p_perm_ = 0.001, spatial-autocorrelation-preserving permutation test). C) Connectivity dimensionality mirrors the U-shaped pattern. *Top:* Brain maps showing connectivity dimensionality (DFC, participation ratio of source-to-target connectivity singular value spectrum) for each region as target. Color scale ranges from 8.91 to 38.16. *Bottom:* Connectivity dimensionality plotted along the sensory-motor gradient, quadratic regression fit in black curve, bootstrapped 95% confidence intervals (grey error band) widen due to fewer regions. While model comparison favors a linear fit (ΔAIC = −9.76), visual inspection shows curvature driven by regions at gradient extremes (black curve: quadratic fit with 95% bootstrap CI). D) Compression and expansion relative to source regions follow the gradient pattern. *Top:* Brain maps showing change in representational dimensionality (ΔDRep = target dimensionality minus average source dimensionality). Negative values (blue, compression) indicate the target has lower dimensionality than its sources; positive values (red, expansion) indicate higher dimensionality than sources. *Bottom:* Compression/expansion plotted along the sensory-motor gradient, recapitulating the U-shaped pattern (quadratic regression fit in black curve, bootstrapped 95% confidence intervals as grey error band). **E) Connectivity dimensionality predicts compression and expansion across all regions.** Direct correlation between connectivity dimensionality and representational compression/expansion across all 360 cortical regions (Pearson Correlation r = 0.44, p < 0.001 using spatial-autocorrelation preserving permutation tests (using BrainSMASH, two-sided); relationship remains significant after controlling for region size, signal-to-noise ratio, inter-vertex distance, temporal variance, and spatial homogeneity). Each dot represents one region, colored by network identity; linear regression fit in black curve, bootstrapped 95% confidence intervals as grey error band. Higher connectivity dimensionality (diverse connectivity patterns among target vertices) predicts representational expansion; lower connectivity dimensionality (similar patterns) predicts compression. (End of Figure 2 caption)

Consistent with our hypothesis, connectivity dimensionality mirrored this trend with a similar U-shaped trajectory (connectivity dimensionality favored a linear fit overall over quadratic (ΔAIC = −9.76) although with visible curvature at gradient extremes). These observations revealed a significant positive relationship between compression and connectivity dimensionality (r = 0.44; group t-test against zero: t(17) = 16.94, p = 4.44e-12; SA-preserving permutation test: p_perm_ < 0.001; Fig. 2E). This positive relationship was consistent across sensory, association, and motor systems (all p < 0.001, Extended Data Fig. 2), demonstrating that this relationship holds across all major cortical systems.

This relationship between connectivity and representational dimensionality is non-trivial for several reasons. First, the two dimensionality measures capture fundamentally different aspects of brain organization. Connectivity dimensionality reflects the geometric structure of how regions connect during rest—essentially, whether vertices of two regions share similar or diverse connectivity patterns. In contrast, representational dimensionality reflects how a single region’s activity varies across 96 different task conditions during active task performance. These are distinct sources of variance measured in different brain states. Second, if representational dimensionality were driven by local (within-region) computations alone, it would be independent of task-independent connectivity properties like cross-region connectivity dimensionality. Third, low-dimensional connectivity could theoretically support representational expansion through non-linear transformations in the target region (e.g., if target neurons applied complex, task-dependent computations to their inputs beyond simple weighted summation). Finally, high-dimensional connectivity might have simply reflected noisy or random connectivity patterns, bearing no relationship to representational dimensionality. To rule out confounds, we regressed anatomical (region size, inter-vertex distance), rest-related (tSNR, signal homogeneity), and task-related (condition reliability, betaSNR) properties from connectivity and representational dimensionality (Extended Data Fig. 3). The compression-expansion relationship remained essentially unchanged (r = 0.4357, p_perm_ < 0.001). To demonstrate that this result is not a given, we constructed a simulation showing that strong non-linear local computations in target regions can decouple the relationship between connectivity dimensionality and representational change (Extended Data Fig. 4), demonstrating that the observed relationship is not mathematically inevitable. Together, these results support our hypothesis that lower-dimensional FC is associated with greater representational compression.

A potential concern is that BOLD signal in association cortex, where mixed-selective neurons are spatially intermingled, may poorly reflect underlying representational diversity^29^. However, we found robust task decodability throughout cortex (Extended Data Fig. 5), with frontoparietal regions comparable to sensory and motor regions; compression was instead concentrated in default-mode and auditory networks, indicating specific functional network organization rather than a general BOLD integration artifact.

### Cross-region activity flows transform representational geometries

While these analyses revealed a robust correlation between connectivity dimensionality and representational compression, they leave the generative question open: does low-dimensional connectivity *produce* representational compression, or merely correlate with it? We addressed this using activity flow modeling^21,22^, which tests whether simulated propagation of task activity over empirical connectivity patterns can reproduce observed representational transformations. This involved comparing model-generated RSMs (by first generating task activation patterns for all task conditions) with those of the true RSMs in target regions. The model-generated estimates were based solely on empirical source activity and task-free source-to-target FC. Strong correspondence between model-generated and empirical representational properties would suggest that connectivity patterns play an important role in shaping representational transformations. Importantly, our goal is not to claim that activity flow provides a complete model of representational transformation. Rather, we test whether connectivity geometry plays a systematic role in shaping these transformations. The linear activity flow model establishes a baseline: if connectivity patterns alone (without additional factors like local circuit dynamics, attention, or neuromodulation) can predict substantial variance in target representations, this demonstrates that connectivity architecture provides fundamental constraints on how representations transform between regions.

Activity flow modeling simulates how task-evoked activity propagates through the network: for each target vertex, we multiply source vertex activity patterns by their connectivity weights and sum the weighted contributions (Fig. 1C). This generates predicted activity patterns in each target region based solely on: (1) observed task activity in source regions, and (2) resting-state connectivity structure. We then compared these model-generated predictions with observed task activity. The model-generated activity patterns were significantly predictive in all regions. Pearson correlations between observed and predicted activity patterns (averaged across subjects, task conditions, and sessions) ranged from r = 0.16 to 0.83 across regions (FDR-corrected p-values: 6.17e-14 to 0.041; Fig. 3A *top*). To robustly test the importance of structured FC patterns in generating the observed activity patterns, we randomly permuted the FC weights between source and target vertices and generated 1000 group-level null means. The observed group-level means were found to be significantly outside the null distribution (p_connperm_ < 0.001 for all target regions, Fig. 3A *bottom*).

**Figure 3:**
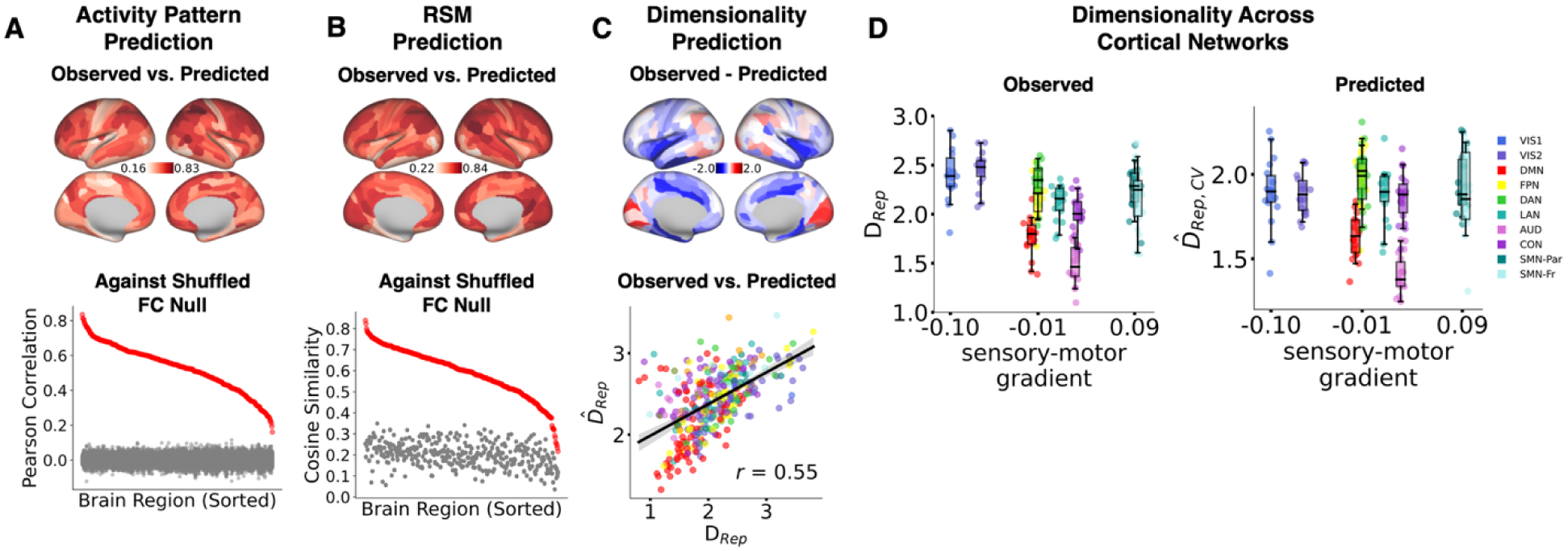
Activity flow over connectivity generates observed representational geometries. A) Activity flow accurately predicts task-evoked activity patterns. *Top:* Brain maps showing Pearson correlation between observed and activity flow-predicted activity patterns for each region, averaged across subjects, sessions, and 96 task conditions. *Bottom:* Prediction accuracy (red) for each region plotted against null distribution (gray) generated by randomly shuffling source-to-target connectivity weights (1000 iterations). All 360 regions show significantly better predictions from intact connectivity than shuffled connectivity (all p < 0.001, two-sided permutation test). A more conservative permutation shuffling only source-side connections, which preserves the target region’s own connectivity structure, confirmed this specificity in 337 of 360 regions (p_connperm_ < 0.05, two-sided permutation test). B) Activity flow captures representational similarity structure. *Top:* Brain maps showing cosine similarity between observed and predicted representational similarity matrices (RSMs) for each region, averaged across subjects. This tests whether connectivity-based predictions capture not just mean activity levels but the full geometry of how task conditions relate to one another. *Bottom:* RSM prediction accuracy (red) significantly exceeds shuffled-FC null (gray) in all regions (all p < 0.001, two-sided permutation test). C) Activity flow predicts representational dimensionality and reveals where additional factors operate. *Top:* Brain maps showing prediction residuals (observed minus predicted representational dimensionality). Blue regions indicate systematic underprediction in early visual cortex and frontoparietal regions, suggesting local dynamics or task-dependent reconfigurations increase dimensionality beyond connectivity-based flow. Red regions indicate overprediction in late visual areas, medial temporal lobe, and salience network, suggesting specialized local computations compress representations beyond connectivity constraints. *Bottom:* Pearson correlation between observed and predicted dimensionality across all 360 regions (r = 0.55, p < 0.001 using spatial-autocorrelation preserving permutation tests, two-sided). Each dot represents one region, colored by network identity; linear regression fit in black curve, bootstrapped 95% confidence intervals as grey error band. The dimensionality prediction was consistent within each cortical system (sensory: r = 0.76, association: r = 0.53, motor: r = 0.66, all p < 0.0001). **D) Activity flow reproduces compression-then-expansion pattern at the subject level.** Representational dimensionality for observed (left) and activity flow-predicted (right) representations, plotted for major cortical networks arranged along the sensory-motor gradient. Each dot represents one subject (n=18); box plots show median and quartiles as box center and bounds, with whiskers indicating the range of datapoints within 1.5 times inter-quartile range. For both observed and predicted data, dimensionality is averaged across all regions within each network for each subject individually, then plotted here. Statistical comparisons (two-sided Paired t-tests, Bonferroni corrected) between networks (VIS1 vs. DMN; DMN vs. SMN-Fr) test whether the compression-then-expansion pattern holds at the individual subject level. The model successfully reproduces the U-shaped pattern: high dimensionality in visual networks, compression in default-mode network, and re-expansion in motor regions. (End of Figure 3 caption)

A more conservative test, shuffling connectivity only on the source side while preserving each target’s intrinsic connectivity structure, confirmed this specificity: 337 of 360 regions still showed significantly better predictions from intact than source-permuted connectivity (p_connperm_ < 0.05), indicating genuine source-to-target structure rather than target-geometry artifacts (Fig. 3A).

We next tested whether these connectivity-generated activity patterns reproduced the observed representational geometry—specifically, the similarity structure across task conditions captured by RSMs. This similarity structure can reveal important patterns, such as how representations generalize across tasks. We tested the ability of observed connectivity patterns to transform representational geometry between regions, as quantified by RSMs and dimensionality. As expected, model-generated and observed RSMs showed strong correspondence across brain regions. Cosine similarity (averaged across subjects) ranged across regions from 0.22 to 0.84, while p_FDR_ values for t-tests against zero ranged from 3.30e-13 to 2.51e-05 (Fig. 3B *top*). Comparison against null values of RSM predictions generated from FC shuffling resulted in group means of all regions’ predictions lying significantly outside of the null distribution (all p_connperm_ < 0.001; source-side shuffling only: p_connperm_ < 0.05 in 358 out of 360 regions; Fig. 3B *bottom*). The model-predicted representational dimensionality values (averaged across subjects) also showed significant correspondence with observed dimensionality values in target regions (r = 0.55, t-test against zero: t(17) = 10.94, p = 4.07e-09, Fig. 3C *bottom*; comparison to FC-shuffled null distribution: p_connperm_ < 0.001; source-side shuffling only: p_connperm_ < 0.001). Predictions were also consistent within each cortical system (sensory: r = 0.76, association: r = 0.53, motor: r = 0.66, all p <0.0001).

Prediction residuals (observed minus predicted dimensionality) revealed systematic under- and overprediction in specific regions (Fig. 3C top), indicating where local circuit dynamics operate beyond the connectivity-based baseline.

We hypothesized that activity flows over observed connectivity patterns also generate the previously observed compression-then-expansion pattern in dimensionality^6^ (Fig. 2B). This pattern of representational transformations is likely essential for flexible performance of diverse tasks, as sensory information is compressed (generalized) across task conditions for learning transfer, before task-specific representational expansions specify task-appropriate motor responses. While we used the same dataset as the original study reporting the compression-then-expansion pattern ^6^, we expanded the number of modeled task conditions from 45 to 96 to better characterize representational dimensionality (see Methods). With this expanded task condition set, we first replicated this compression-then-expansion pattern in our data’s task condition split. Examining network-level dimensionality patterns, we found that visual (VIS1) and motor (SMN-Fr) networks showed significantly higher dimensionality than default-mode (DMN) and auditory (AUD) networks, which exhibited the strongest compression (VIS1 vs. DMN: t(17) = 9.81, p = 2.05e-08; SMN-Fr vs. DMN: t(17) = 6.46, p = 5.86e-06; VIS1 vs. AUD: t(17) = 13.61, p = 1.43e-10; SMN-Fr vs. AUD: t(17) = 10.02, p = 1.49e-08; Fig. 3D *left*). To test whether the activity flow model could reproduce this pattern, we employed a double cross-validated approach (see Methods): we constructed hybrid RSMs by measuring the cosine similarity between observed activity patterns of tasks from one run and predicted activity patterns of tasks from a different run. We then analyzed the dimensionality of these double cross-validated RSMs using the participation ratio measure. Our analysis revealed that the model successfully reconstructed the compression-then-expansion pattern of dimensionality along the sensory-association-motor axis, with visual and motor networks again showing significantly higher dimensionality than default-mode and auditory networks (VIS1 vs. DMN: t(17) = 5.49, p = 4.02e-05; SMN-Fr vs. DMN: t(17) = 5.11, p = 8.65e-05; VIS1 vs. AUD: t(17) = 9.57, p = 2.93e-08; SMN-Fr vs. AUD: t(17) = 13.93, p = 1.00e-10; Fig. 3D *right*). Together, these results demonstrate that activity flows shaped by specific geometric properties of connectivity patterns generate the observed representational geometries.

### Activity flows over connectivity patterns drive transformations of task representations

Previous analyses established that empirical connectivity patterns reliably generate observed representational geometries in target regions. However, it remains possible that the representations generated by the activity flow models transformed representations only minimally, with similarities between source and target representations explaining the accurate predictions of target representations. For instance, a source and a target region may both represent the word “green” during the Stroop paradigm, with the activity flow model simply transferring (rather than truly transforming) a representation between these two regions. This would result in flow-predicted RSMs that are no more similar to the target than the source. Further, it remains possible that predicted RSMs of a target region were more similar to the source RSM than the true target RSM. Instead, we hypothesized that activity flows implement true representational transformations most of the time, predicting that generated RSMs would be more similar to the target RSM than the source RSM for most regions. We therefore next examined whether the geometry of connectivity-generated predictions more closely resembled their source or target geometries.

We quantified this using two cosine distances: d_pred_ (predicted-to-target RSM) and d _transform_(predicted-to-source RSM); d_pred_ < d _transform_ indicates genuine transformation rather than transfer (Extended Data Fig. 6A). Connectivity-based transformations were non-random, with 357 of 360 regions differing significantly from shuffled-connectivity nulls (p_connperm_ < 0.001; Extended Data Fig. 6C), and predicted transformation magnitude closely tracked observed source-to-target distances (r = 0.62, p_perm_ < 0.001; Extended Data Fig. 6B).

Predicted geometries more closely matched target than source geometries in 339 of 360 regions (p_FDR_ < 0.05; 292 at p_FDR_ < 0.0001; Extended Data Fig. 6D), supporting a fundamental role for connectivity patterns in transforming representational geometries across regions. The regions where predictions were closest to targets (smallest d_pred_) versus sources included both compression zones (association cortex, where low-dimensional connectivity compressed representations) and expansion zones (motor cortex, where high-dimensional connectivity expanded representations), demonstrating that connectivity geometry shapes transformations bidirectionally.

### Low-dimensional connectivity shapes cross-task representational similarity

The transition from sensory to association cortex was accompanied by substantial reductions in FC dimensionality (Fig. 2C). To explore the cognitive benefits of such low-dimensional connectivity, we tested whether target regions with lower connectivity dimensionality show greater similarity among task representations, reflecting latent factors shared across tasks that support the brain’s capacity for diverse task performance.

To isolate cross-task similarity while excluding within-task effects, we computed mean cosine similarities between task condition combinations for each task pair (16 tasks), retaining only positive values across subjects (Fig. 4A, see Methods).

**Figure 4:**
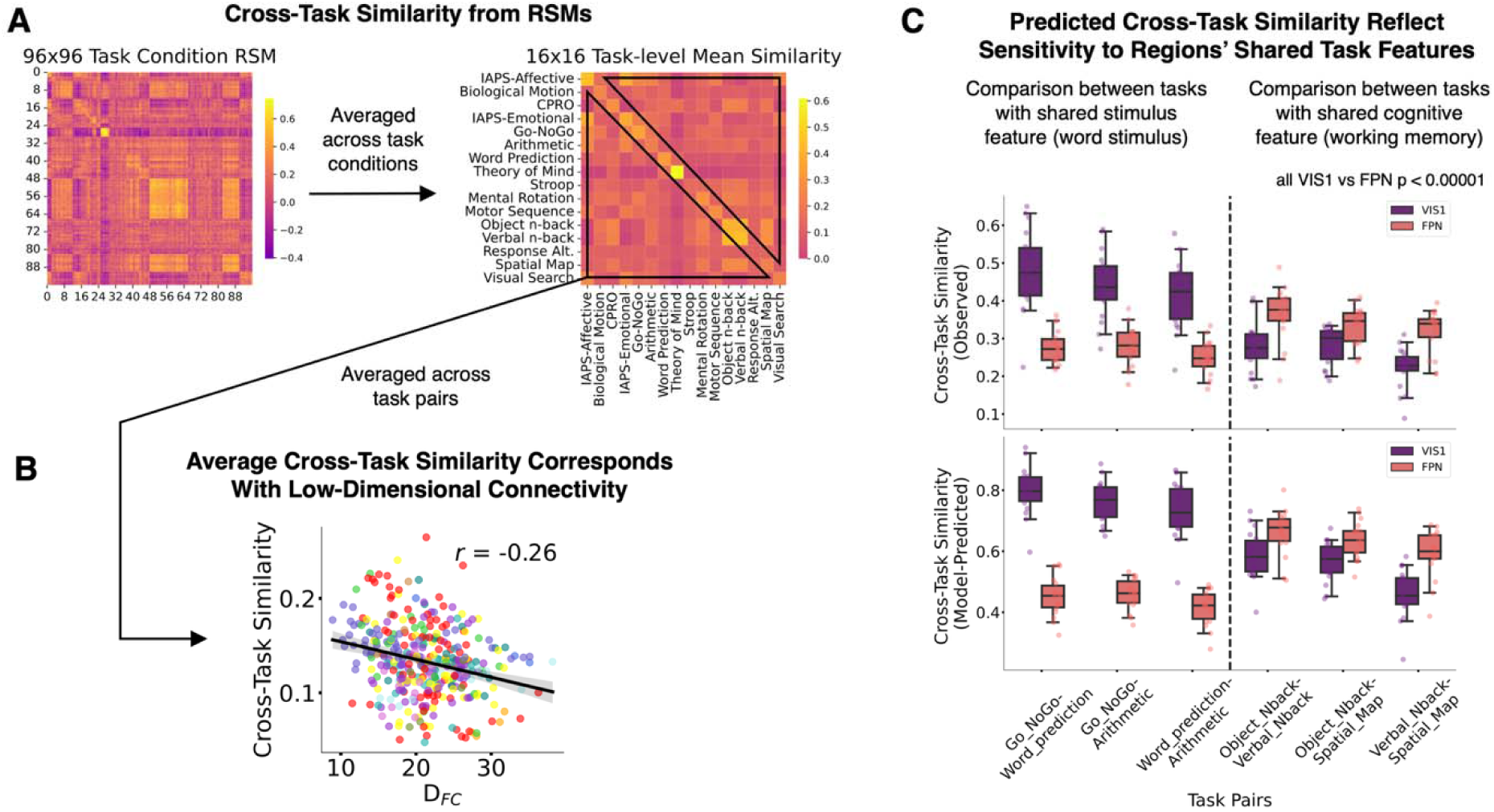
Low-dimensional connectivity generates functionally interpretable cross-task similarities. A) Deriving cross-task similarity from representational similarity matrices. Cross-task similarity quantifies how similar a region’s representations are across different tasks, revealing shared latent factors. *Left:* Example 96×96 task condition RSM from a single region, showing cosine similarity between all pairs of task conditions (averaged across subjects). Blocked structure reflects tasks with multiple conditions (e.g., conditions 1-8 are Stroop task conditions, showing high within-task similarity). *Right:* To isolate cross-task similarity, we first retained only positive similarity values from the 96×96 condition RSM (negative similarities are difficult to interpret), then averaged these positive values within each task pair to create a 16×16 task-level matrix. Off-diagonal elements represent cross-task similarity (e.g., how similar are representations between Stroop and Word Prediction tasks?). For each region, we computed mean cross-task similarity by averaging all off-diagonal values in this 16×16 matrix, yielding a single measure of how much the region shares representational structure across tasks. **B) Regions with low-dimensional connectivity show higher cross-task similarity.** Pearson correlation between connectivity dimensionality (DFC) and mean cross-task similarity across all 360 regions (r = −0.26, p < 0.001 using spatial-autocorrelation preserving permutation tests, two-sided). Each dot represents one region, colored by network identity; linear fit in black curve, bootstrapped 95% confidence intervals as grey error band. Lower connectivity dimensionality (similar connectivity patterns among target vertices) predicts greater similarity of task representations, suggesting these regions encode shared latent factors that generalize across tasks. Higher connectivity dimensionality predicts task-specific representations. **C) Activity flow-predicted cross-task similarities reflect network-specific functional specialization.** Comparison of cross-task similarity patterns between primary visual network (VIS1, purple) and frontoparietal network (FPN, red) for task pairs with shared features. For each subject, cross-task similarity values were averaged across all regions within each network. Each box plot shows distribution across subjects (n=18); each dot represents one subject’s network-averaged cross-task similarity for that task pair; box plots show median and quartiles as box center and bounds, with whiskers indicating the range of datapoints within 1.5 times inter-quartile range. *Top:* Observed cross-task similarities. *Bottom:* Model-predicted cross-task similarities from activity flow. *Left panel:* Task pairs sharing sensory features (word stimuli: *Go-NoGo*, *Word Prediction*, and *Arithmetic* tasks): VIS1 shows significantly higher cross-task similarity than FPN across all three task pairs shown (all p < 0.00001, two-sided paired t-tests, Bonferroni corrected, exact values of all comparisons in Extended Data), reflecting VIS1’s sensitivity to shared visual word processing demands. *Right panel:* Task pairs sharing cognitive features (working memory: *Object n-back*, *Verbal n-back*, and *Spatial Map* tasks): FPN shows higher cross-task similarity than VIS1, reflecting FPN’s sensitivity to shared cognitive control demands. Critically, activity flow predictions (bottom) reproduce these network-specific patterns, demonstrating that connectivity geometry alone generates functionally interpretable latent task factors aligned with each network’s computational specialization. (End of Figure 4 caption)

Our analysis yielded two key findings. First, we hypothesized that low-dimensional connectivity actively *generates* cross-task similarities—not merely that low-dimensionality regions happen to show similar representations, but that connectivity geometry creates these similarities through activity flow. Consistent with this, regions with lower-dimensional connectivity showed stronger cross-task representational similarity across all task pairs (r = −0.26, t(17) = −24.92, p = 8.01e-15; SA-preserving permutation test: p_perm_ = 0.01; Fig. 4B).

Activity flow over these same connectivity patterns reproduced the observed cross-task similarity structure across most regions (cosine similarity 0.13–0.99; p_connperm_ < 0.05 in 355/360 regions; Extended Data Fig. 7A).

A simulation using random source activity (ensuring near-zero cross-task similarity at the source) confirmed this as a general property of connectivity-based transformations: lower-dimensional connectivity consistently produced greater cross-task similarity (ρ = −0.988, p = 1.08e-72; see Methods).

To begin exploring whether these cross-task similarities might reflect interpretable task factors, we conducted an exploratory analysis at the network level by averaging the cross-task similarity patterns of regions among twelve cortex-wide FC networks according to an established network parcellation^30^. While our task set was not designed to systematically investigate shared cognitive components, selective examination of two networks revealed potentially meaningful contrasts. For instance, in the primary visual network (VIS1), we observed high cross-task similarity among the *Go-NoGo*, *Word Prediction* and *Arithmetic* tasks, which share visual word processing demands. In contrast, the fronto-parietal network (FPN) showed high similarity between the *Object n-back*, *Verbal n-back* and *Spatial Map* tasks despite their different stimulus types, likely reflecting shared cognitive control demands (Fig. 4C *top*).

This pattern of network-relevant latent task factors was replicated when connectivity-predicted estimates were analyzed, demonstrating the role of connectivity patterns in generation of meaningful latent task factors (Fig. 4C *bottom*). These network-specific patterns are non-trivial: VIS1’s high similarity for word-processing tasks and FPN’s high similarity for cognitive control tasks demonstrate that connectivity geometry generates functionally interpretable latent task factors aligned with each network’s computational role (see observed and predicted network comparison results in Extended Data Table 1). Cross-task similarities at the network level significantly differed from null models based on shuffled connectivity, for all networks (Extended Data Fig. 7B). Future work with a more systematic task battery, perhaps involving reuse of instruction elements across tasks (e.g., to allow for cross-condition generalization analysis), would be needed to thoroughly investigate these network- and region-specific patterns.

Together, these results show that low-dimensional connectivity generates shared representational structure across tasks, further supporting a fundamental role for intrinsic connectivity in specifying task representations throughout cortex.

### Activity flow-driven expansions generate task-variable conjunctions

We hypothesized that connectivity patterns expanding representational dimensionality specifically support task-variable conjunctions: activity patterns present only when two or more task features co-occur (e.g., ink color blue *and* Stroop-incongruent), rather than whenever any single feature is present. Such conjunctive coding allows cognitive variables to interact in situation-specific ways—for instance, processing color blue differently depending on congruency—unlike low-dimensional compositional representations that are reused identically across contexts. While our task set was not systematically designed to investigate conjunctive coding, we identified a subset of five tasks containing separable sensorimotor and cognitive variables that allowed for an exploratory test of this hypothesis.

To test which task features drive representational similarity, we identified five tasks with separable sensorimotor and cognitive variables (Fig. 5A) and constructed four template RSMs: (1) sensory similarity (based on stimulus features like color, shape, words), (2) cognitive similarity (based on task demands like congruency, difficulty), (3) motor similarity (based on required button press responses), and (4) conjunction similarity (multiplicative combination of sensory × cognitive × motor features, capturing non-additive interactions). Using ridge regression, we fit these four vectorized matrices to the vectorized observed neural RSM (Fig. 5A).

**Figure 5:**
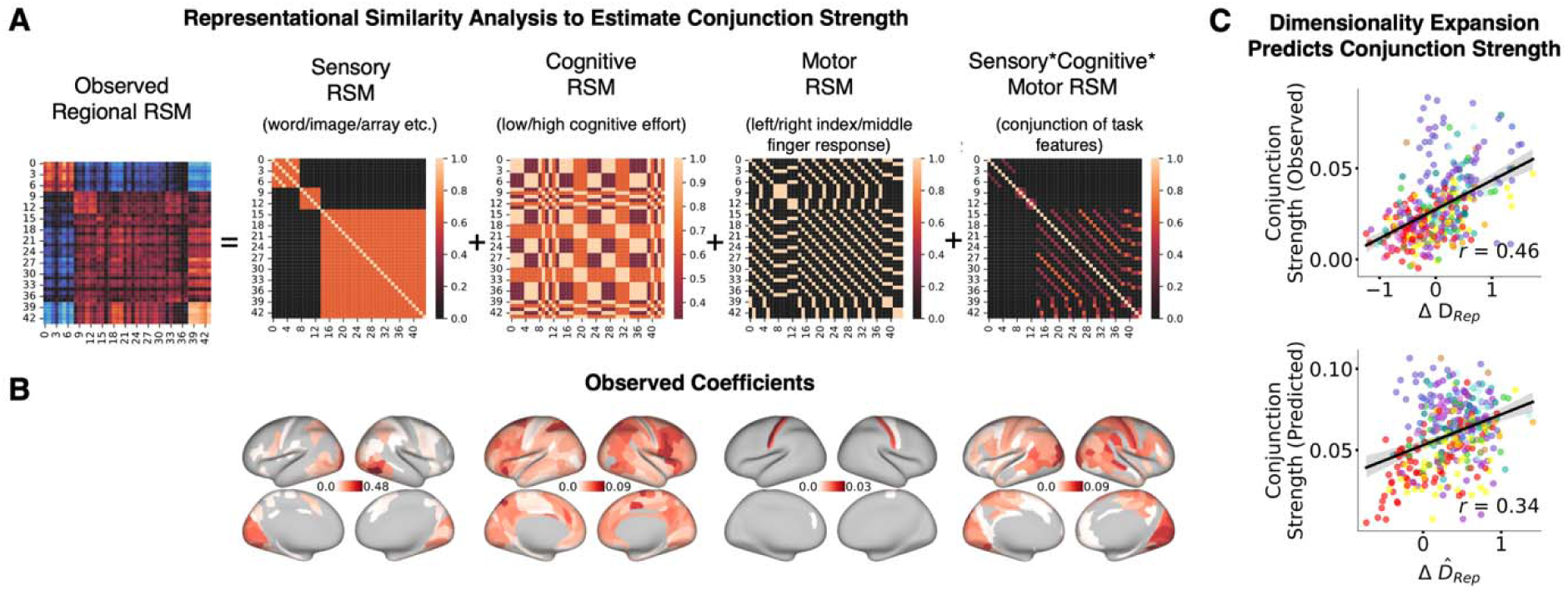
Connectivity-driven dimensionality expansion supports conjunctive encoding of task features. A) Representational similarity analysis to estimate conjunction strength. We identified a subset of 5 tasks (44 task conditions total) with separable sensorimotor and cognitive features: Stroop, Go/NoGo, IAPS-Affective, IAPS-Emotional, and Mental Rotation. For each region, we fit the observed 44×44 task condition RSM (left; example shown from right Area PH in visual cortex) using ridge regression against four template RSMs encoding different feature types. *Sensory template:* similarity based on stimulus type (word, image, spatial array). *Cognitive template:* similarity based on task demand level (low vs. high cognitive effort). *Motor template:* similarity based on required button press responses (left/right index/middle finger). *Conjunction template:* multiplicative combination of all three features (sensory × cognitive × motor), capturing non-additive feature interactions where specific combinations (e.g., “word stimulus + high demand + left index response”) produce distinct representations beyond the sum of individual features. Ridge regression yields four coefficients per region, with the conjunction coefficient quantifying how strongly that region encodes feature combinations. **B) Observed feature encoding shows expected anatomical specificity.** Brain maps showing fitted coefficients (averaged across subjects) for each feature template, with only significant values displayed (one-sample t-test against zero, FDR corrected). *Sensory:* highest in visual cortex (9/10 highest coefficients). *Cognitive:* highest in association cortex (7/10 highest in frontoparietal and default-mode regions). *Motor:* highest in motor and premotor cortex (8/10 highest). *Conjunction:* more distributed, with highest values in regions showing representational expansion. This anatomical specificity validates the RSA approach and demonstrates that sensory, cognitive, and motor features are encoded in their respective brain systems. **C) Dimensionality expansion predicts conjunction strength in both observed and predicted data.** *Top:* Correlation between observed conjunction coefficient and observed representational compression/expansion (ΔD_Rep_) across all 360 regions (r = 0.46, p < 0.001 using spatial-autocorrelation preserving permutation tests, two-sided; linear fit in black curve, bootstrapped 95% confidence intervals as grey error band). Each dot represents one region, colored by network. Regions showing expansion (positive ΔD_Rep_) encode stronger conjunctive representations, while regions showing compression encode weaker conjunctions. This relationship is stronger than correlations with individual feature terms (sensory: r = 0.15; cognitive: r = 0.28; motor: r = 0.04), demonstrating that expansion specifically supports conjunctive coding. *Bottom:* Correlation between activity flow-predicted conjunction coefficient and predicted ΔD_Rep_ (r = 0.34, p < 0.001 using spatial-autocorrelation preserving permutation tests, two-sided; linear fit in black curve, bootstrapped 95% confidence intervals as grey error band). The model reproduces the relationship between expansion and conjunction strength using only connectivity geometry and source activity, demonstrating that high-dimensional connectivity patterns systematically support conjunctive representations. (End of Figure 5 caption)

As expected, sensory, cognitive, and motor similarities were encoded in their respective brain systems (Fig. 5B). Critically, conjunction-term coefficients correlated positively with representational expansion (r_conj_ = 0.46, p = 2.92e-07; Fig. 5C top), a relationship stronger than for cognitive or motor terms individually and comparable to the sensory term (Extended Data Table 2). Activity flow-predicted conjunction strength similarly correlated with predicted expansion (r = 0.34, p = 0.006; Fig. 5C bottom), exceeding predicted correlations for the other three terms (Extended Data Table 2), confirming that connectivity-driven expansion specifically supports conjunctive coding.

Beyond conjunctive coding, we also tested whether representational dimensionality relates to cognitive flexibility more broadly, using fluid intelligence data from Human Connectome Project^27,28^ (HCP; n=352). Region-averaged dimensionality positively predicted general fluid intelligence (r=0.29, p=1.5e-08), particularly in sensory and association regions (Extended Data Fig. 8 A-D). These findings generate specific hypotheses that can be tested in future studies using carefully designed task batteries that systematically vary sensory, cognitive, and motor demands to more definitively determine how connectivity-based transformations contribute to high-dimensional task representations and support flexible cognitive control through conjunctive coding. As a replication check, we checked for generalizability of the compression-then-expansion pattern in representational dimensionality in the HCP dataset as well. Using cross-validated RSMs constructed from 24 task conditions across seven tasks, we found that regional representational dimensionality exhibited the same U-shaped pattern along the sensorimotor gradient at the group level (Extended Data Fig. 8E). To quantify the consistency of compression in association regions relative to sensory and motor regions across individuals, we calculated a compression ratio for each subject (mean dimensionality of sensory and motor regions divided by association region dimensionality). A compression ratio greater than 1 indicated representational compression in association cortices. This pattern proved highly reliable, with 98.6% (347/352) of subjects showing compression in association regions, providing strong validation of this organizational principle (Extended Data Fig. 8F).

## Discussion

We identified a systems-level organizing principle by which intrinsic connectivity dimensionality directs the cross-region compression and expansion of task representations: low-dimensional connectivity produces compressed representations with shared similarities across tasks, while high-dimensional connectivity expands representations to integrate task variables for specific cognitive demands. Activity flow modeling demonstrated that this connectivity-based mechanism actively generates observed representational geometries across sensory, association, and motor systems, rather than merely correlating with them. Regions with lower-dimensional connectivity showed stronger cross-task similarities, suggesting shared latent features supporting generalization, while high-dimensional regions supported conjunctive representations situating that information for specific contexts. This mechanism replicated in an independent HCP dataset, where representational dimensionality also predicted general fluid intelligence.

Our simulations show that strong non-linear local computations can fully decouple connectivity dimensionality from representational transformations (Extended Data Fig. 4), yet we empirically observe robust coupling (r=0.44), positioning the brain in an intermediate regime where network architecture provides systematic constraints while local circuit dynamics contribute region-specific modulation, as evidenced by our prediction residuals (Fig. 3C). This suggests a design principle of architectural parsimony: connectivity patterns, once established, provide stable constraints shaping representations across tasks without requiring task-dependent reconfiguration of local circuits, with residual patterns indicating a principled partitioning of computational labor between distributed architecture and specialized local dynamics. This framework generates testable predictions: diseases affecting white matter connectivity should produce predictable distortions in representational geometry, and connectivity-targeted interventions should have systematic downstream effects on representational transformations.

Our findings provide empirical validation of theoretical predictions from computational neuroscience regarding how connectivity structure constrains representational geometry^18–20,31–33^, which have been extensively characterized in artificial neural networks, but not previously demonstrated in biological circuits ^34^. Our resting-state FC-based approach offers a functionally defined measure of connectivity constraints complementary to the structural connectivity emphasized in theoretical work.

The present study expands previous connectivity-based models of task processing from structured rule combinations to diverse cognitive demands. Earlier work using the Concrete Permuted Rules Operation (C-PRO) paradigm demonstrated that task rule information transfers between brain regions via resting-state connectivity ^22^; we show this principle generalizes to diverse cognitive tasks lacking such structured relationships, extending the activity flow modeling framework to show connectivity patterns actively transform, not merely transfer, task representations between regions. While previous work emphasized information transfer between regions encoding similar information, we demonstrate that intrinsic connectivity patterns support representational transformations between functionally diverse connected regions. Our results also extend prior findings showing that pairwise similarity between regional RSMs correlates with resting-state connectivity at the parcel level ^6^. We build upon this finding by developing a mechanistic model based on finer-grained, vertex-level connectivity that characterizes how connectivity geometry constrains these representational transformations..

Our study instantiates a key theoretical framework for understanding cognitive flexibility: examining how multiple tasks mutually constrain neural activity patterns to reveal mechanisms of generalization ^35^. Activity flows over intrinsic connectivity generated widespread shared representations across 355 of 360 regions, comparable to cognitive encoding models explaining variance in 86% of cortical voxels ^2^, but through direct connectivity-mediated mappings rather than predefined cognitive factors ^5^. This connectivity-based framework leverages a brain-based ontology^36^ that remains open to discovering latent cross-task factors not corresponding to existing cognitive constructs.

Our findings complement a growing body of work in animal models examining how inter-regional connectivity shapes neural computation^37–39^. Studies in non-human primates have demonstrated that functional connectivity between sensory and association areas constrains how information is communicated and transformed across regions^40^, with communication subspaces between areas shaping what information can be transmitted^41–43^. Recent work in mice similarly shows that inter-regional coordination structures cortical dynamics and supports flexible behavior^44^. Our findings extend these principles to human whole-brain organization during diverse cognitive tasks, demonstrating that connectivity geometry operates as a systematic organizing principle across species and scales, from sensory processing in animal models to cognitive flexibility in humans. A remaining limitation is that BOLD-based measures cannot resolve contributions of mixed selectivity at the sub-voxel level, which may manifest as low dimensionality and low decodability and could also affect connectivity dimensionality estimation. Resolving this fully requires cross-modal validation — for example, estimating connectivity dimensionality from noise correlations between neuronal populations in electrophysiology recordings, and testing whether this predicts representational dimensionality change between those populations across task conditions.

A key conceptual clarification: activity flow modeling here involves literal convergence (many source vertices projecting to fewer target vertices), yet we define compression/expansion relative to the average dimensionality of individual source regions rather than pooled source-vertex dimensionality, allowing us to characterize region-to-region transformations while acknowledging the underlying integration across many sources. This framework reveals that convergent architectures can produce dimensionality expansion through a plausible linear mechanism: when source regions occupy distinct low-dimensional subspaces and target vertices receive diverse weighted mixtures (high connectivity dimensionality), the target can occupy a higher-dimensional space spanning combinations of those subspaces—potentially explaining our conjunctive coding results (Fig. 5), since linear mixing of diverse sources naturally creates feature combinations approximating conjunctive coding. This generates a testable prediction: expansion-showing regions should receive input from sources with lower cross-source representational similarity.

The present results reveal how the brain’s intrinsic connectivity architecture enables both generalization across tasks and specific task implementation through systematic control of representational dimensionality. The geometric properties of neural representations have proven highly informative for understanding how brain regions support multiple task contexts ^8,11,14,17^. Building on this framework, we show that these geometries are fundamentally shaped by connectivity patterns: low-dimensional connectivity supports abstraction of shared features across tasks, as evidenced by meaningful task groupings in our network-level analyses – such as visual word processing demands in visual networks and cognitive control demands in frontoparietal networks. These findings align with the neural reuse framework^45,46^, which proposes that brain regions and circuits support multiple cognitive functions by sharing computational resources across tasks; here, we characterize this reuse in terms of the geometric properties of connectivity and representations. Complementarily, high-dimensional connectivity enables the integration of multiple task variables into conjunctive representations, which we measured using established RSA methods ^15,16^ to show how abstract features are situated for specific cognitive demands. For example, a task-implementing conjunctive representation may only be active when the rule representation “press your left index finger when you see green” is active and a representation of green is active, with activity flows from that representation to a left index finger representation in the motor system implementing task behavior. This connectivity-based mechanism – abstraction through compression and specificity through expansion – provides a principled basis for cognitive flexibility, enabling both generalization across tasks and task-specific processing.

Higher representational dimensionality, particularly in sensory and association regions, predicted general fluid intelligence^47^—a finding that appears to conflict with the compression-then-expansion pattern, but likely reflects complementary levels of organization: compression-then-expansion is a canonical within-subject architecture supporting efficient task processing, while absolute dimensionality within that architecture reflects individual capacity for flexible cognition (Extended Data Fig. S8).

In summary, our study reveals a fundamental organizing principle by which intrinsic connectivity architecture enables cognitive flexibility through systematic transformation of task representations, providing a framework for understanding both healthy cognitive function and its disruption in neurological and psychiatric conditions.

## Supporting information

Inventory of Supporting Information

## Acknowledgements

Multi-domain Task Battery data were provided by Maedbh King, Carlos Hernandez-Castillo, Russell A. Poldrack, Jonathon Walters, Richard B. Ivry, Jörn Diedrichsen via OpenNeuro (doi:10.18112/openneuro.ds002105.v1.1.0). HCP data were provided by the Human Connectome Project, WU-Minn Consortium (Principal Investigators: David Van Essen and Kamil Ugurbil; 1U54MH091657) funded by the 16 NIH Institutes and Centers that support the NIH Blueprint for Neuroscience Research; and by the McDonnell Center for Systems Neuroscience at Washington University. We thank the Office of Advanced Research Computing at Rutgers, The State University of New Jersey, for providing access to the Amarel cluster and associated research computing resources that have contributed to the results reported here. The authors thank the members of Cole Neurocognition Lab for helpful conversations during the course of the research.

## Author Contributions Statement

L.N.C., T.I. and M.W.C. conceptualized the project and wrote the paper. L.N.C. performed the formal analysis and visualization, developed software and wrote the original draft. A.T. collaborated in discussions of functional connectivity computation. M.W.C. acquired funding. M.W.C. (senior author) and T.I. supervised the project.

## Methods

### Multi-domain task battery dataset

Portions of this section are paraphrased from the original data study ^48^.

We used the multi-domain task battery (MDTB) dataset ^48^, originally designed to examine functional task boundaries in the human cerebellum. The dataset comprises whole-brain resting-state and task-state fMRI data from 24 right-handed participants (16 females, 8 males; mean age 23.8 ± 2.6 years) at Western University. Participants provided informed consent under an approved experimental protocol. 26 participants were originally scanned over two sessions, but two of them were excluded for not completing all the scanning runs. Participants were compensated based on experimental protocol approved by the institutional review board at Western University, where the data was originally collected.

The MDTB includes 26 cognitive tasks with up to 45 distinct task conditions per participant, divided into sets A and B. Participants completed set A and B tasks in separate identical sessions, with inter-session intervals of 2-3 weeks for half the participants and approximately one year for the others. Each set involved two 10-minute imaging runs. For 18 out of 24 participants, an additional resting-state scan (two 10-minute runs) was conducted separately from the ‘rest’ block in task sessions. Our analyses utilize task data from these 18 participants to relate task and resting-state data within the same individuals.

The dataset encompasses a broad range of cognitive processes. Sets A and B shared eight common tasks and nine unique tasks each, totaling 17 tasks per set. Tasks were performed once per imaging run using an interleaved block design. Each task block consisted of a 5-second instruction screen followed by 30 seconds of continuous task performance, totaling 35-second blocks. Most tasks comprised 10-15 trials per block, with the number of trials per task ranging from 1 to 30. For 17 tasks requiring motor input, participants used a four-button box with index or middle fingers of either or both hands. All tasks within each set were completed in a single imaging run, ensuring a consistent baseline across tasks for all participants.

We selected 16 active, visual tasks (excluding the active auditory interval timing task), decomposing them into 96 task conditions based on stimulus type, motor response, and context variables (Supplementary Data Table 1; decomposition unique to this study). Focusing on visual tasks allowed us to treat visual regions as sensory in the sensory-association-motor hierarchy. fMRI acquisition parameters were as follows: repetition time = 1 s; field-of-view = 20.8 cm; phase encoding direction P to A; 48 slices; 3 mm thickness; in-plane resolution 2.5 × 2.5 mm^2^.

### fMRI preprocessing

This section is paraphrased from the preprocessing details of Ito & Murray (2023) ^6^, which used the same preprocessing pipeline.

The preprocessing of the fMRI data in this study utilized the Human Connectome Project (HCP) preprocessing pipeline^49^, which was executed through the Quantitative Neuroimaging Environment & Toolbox [QuNex, version 0.61.17] ^50^. The HCP pipeline encompassed several key steps in data preparation. These included the reconstruction and segmentation of anatomical images, similar procedures for EPI (echo-planar imaging) data, alignment of brain images to the standardized MNI152 template for spatial normalization, and corrections for head movement during scanning. In addition to the standard HCP pipeline, we applied further steps of nuisance regression, as detailed in the next section.

### fMRI task activation estimation

This study employed a modified version of the single-subject beta series regression technique to analyze the fMRI task data ^51^. The analysis focused on estimating vertex-wise activations throughout the cortical sheet. Each task condition was represented by a distinct task regressor, modeled as boxcar functions (with 1 for on during each trial period of the corresponding task condition and 0 for off, with a separate regressor for each task’s instruction period). Given high practice accuracy of tasks (>85%) prior to scanning, we did not exclude error trials during this estimation. These regressors were then convolved with the SPM canonical hemodynamic response function to account for temporal lags in hemodynamic responses ^52^. The ‘rest’ block in each run was left unmodeled and thus formed part of the baseline. The coefficients of these regressors were used to represent task condition activations.

Task GLMs were fit per run using ridge regression (Python 3.8.5, ’ridge_corr’ function, https://github.com/alexhuth/ridge/blob/master/ridge.py) to address multicollinearity from closely spaced task conditions. In each run, the optimal regularization parameter (alpha) was selected using four-fold cross-validation, testing a range of alpha values from 1 to 1000 in 20 logarithmic steps. The optimal alpha was identified by maximizing the average correlation with the testing fold across cross-validation iterations. Once the optimal alpha was determined, task estimates were obtained for the entire run using this value.

To minimize non-neural noise and avoid potential artifacts, task GLMs were performed concurrently with nuisance regression, as suggested by previous research ^53^. The nuisance parameters included 24 motion-related regressors (six motion parameters, their time-derivatives, and quadratic terms) and eight physiological nuisance signals (mean time series from white matter and ventricle voxels, their derivatives, and quadratic terms). In total, 32 nuisance regressors were incorporated alongside the task regressors to extract refined task activation estimates.

Prior to running the regression, both the time series data and the regressor time series were z-scored along the time axis. Multiple values of activity estimates across different runs were averaged to obtain one estimate per task condition per session. To assess the reliability of activity estimates, we computed Pearson correlation between the estimates from the two identical sessions after averaging across task conditions in each parcel (region) for each subject (range across parcels, averaged across subjects: 0.14 – 0.83). While these correlations were not used to filter out any task conditions, regions, or subjects, they provided a measure of estimated stability across sessions.

### FC estimation

We used the two 10-minute resting-state runs per subject to measure FC. Following preprocessing and nuisance regression (as above), residual timeseries from the two runs were concatenated per subject before estimating FC in two steps.

First, we estimated sparse parcel-level FC between mean Glasser-parcellation ^54^ timeseries using graphical lasso (regularized partial correlation, L1 penalty; ’GGLasso’ package^55^), which improves validity against structural connectivity and individual-level reliability^56^. This two-step approach—sparse parcel-level graph followed by dense vertex-level estimation—balances statistical power with spatial specificity while avoiding the intractability of estimating full vertex-to-vertex connectivity across cortex (millions of connections). The regularization parameter (alpha) was optimized over 0.001–1 (10 log-spaced steps), selecting the value minimizing negative log-likelihood averaged across folds.

The parcel-level graph identified each region’s connected neighbors (sources). Dense vertex-level FC was then estimated between each target region’s vertices and all vertices of its source regions, producing a rectangular connectivity matrix (source vertices × target vertices) per target region. For each target vertex, source vertices served as regressors in a cross-validated principal component regression, chosen to handle thousands of source vertices with only ∼1,200 timepoints while reducing overfitting and preserving major covariance patterns^21,22,57^. The number of components was selected via 10-fold cross-validation (minimizing mean squared error), yielding a mean of 57.1 components across subjects (range: 51–71). Source vertices within 10 mm of the target region’s boundary were excluded to prevent circularity from spatial autocorrelation. Goodness of fit (r²) of the regression to training data ranged from 0.24–0.66 across regions, averaged across subjects.

### RSM estimation

For each of the 360 Glasser parcels, we computed cosine similarity between task condition activity vectors, cross-validated between the two sessions (e.g., condition A from session 1 vs. condition B from session 2). Cosine similarity was chosen over Pearson correlation because it captures both activation pattern and magnitude across vertices. Cross-validating between sessions yielded a non-trivial diagonal, reflecting test-retest reliability. This produced a 96×96 RSM per parcel per subject, corresponding to the 96 task conditions (16 tasks).

### Representational and connectivity dimensionality

We measured representational dimensionality using the participation ratio of the RSM’s eigenvalue spectrum (eigen decomposition), which quantifies the rate of spectral decay:

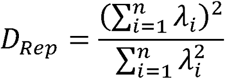

where λ*_i_*are the eigenvalues of the RSM. This measure corresponds to the number of principal components explaining ∼80% of variance^58^, but was preferred for its continuous scale, enabling finer-grained comparisons across regions and subjects.

We related representational dimensionality to decoding of 96 task conditions using a minimum-distance classifier based on cosine distance (cross-validated across two sessions), intuitable directly from the RSM: a condition is decodable if its diagonal value exceeds the other values in its row/column.

For connectivity dimensionality, we sought a parallel measure accounting for the rectangular structure of connectivity data (connecting distinct source and target vertex sets). We performed singular value decomposition on the dense FC matrix between each target region’s vertices and its source regions:

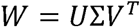

We then calculated the participation ratio of the singular value spectrum as a continuous measure of connectivity dimensionality for each target region:

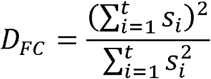

Where *s_i_*are singular values following singular value decomposition.

These measures were calculated separately for each subject. Higher representational dimensionality indicates more distinct condition-specific patterns; lower connectivity dimensionality reflects more convergent, homogeneous source input (see Results for interpretation).

We tested parcel-specific properties that could confound both dimensionality measures: target region size (vertex count), signal-to-noise ratio (mean RSM diagonal), mean inter-vertex distance (mean geodesic distance to neighboring mesh-connected vertices^59^), temporal variance of resting-state activity (SD of each vertex’s time series across time, averaged across vertices), and spatial homogeneity (SD across vertices within each parcel at each timepoint, averaged across time). These confounds were regressed out of connectivity dimensionality prior to correlating the residuals with representational dimensionality. Across-parcel correlations used spatial autocorrelation-preserving permutation tests (BrainSMASH toolbox^60^, 1000 null surrogate maps making the smallest p-value to be 0.001, with any value under that reported as p < 0.001).

### Discrete brain systems and continuous FC gradient estimation

We delineated sensory, association, and motor regions using the Glasser et al. (2016) multi-modal parcellation (HCP-MMP1.0; 180 areas/hemisphere), selected for its integration of cortical architecture (myelin, thickness), function (task-fMRI), connectivity, and topography, providing improved spatial localization over unimodal approaches^54,61^, combined with the Cole-Anticevic Brain Network Partition (CAB-NP)^30^, which divides the parcellation into 12 functional networks. We categorized primary and secondary visual network regions as sensory (restricted to visual networks given our focus on visual tasks); somatomotor regions were split into somatosensory (SMN-Parietal, posterior to the central sulcus) and motor (SMN-Frontal, anterior); all remaining regions were categorized as association.

To derive continuous connectivity gradients, we thresholded the parcel-level sparse FC graph (averaged across subjects) to retain the top 20% of values^62^, then applied principal component analysis. The first component ran from unimodal (sensory/motor) to transmodal (association) regions; the second exhibited continuous variation from sensory through association to motor regions, representing the broad cortical processing hierarchy—we refer to this as the sensorimotor FC gradient, which we visually confirmed replicates prior reports and complements our discrete network categorization.

### Connectivity pattern similarity and Sparsity

To interpret connectivity dimensionality in concrete terms, we analyzed target-vertex connectivity patterns (vectors of connectivity weights from source vertices to each target vertex), motivated by the activity flow framework’s focus on how activity is generated in target populations; connectivity dimensionality is also capped at target region size, the smaller dimension of the dense connectivity matrix. For each target region, we computed the Pearson correlation between all pairs of vertices’ connectivity patterns (t × t matrix, t = number of vertices) and derived the mean correlation as a measure of overall pattern similarity.

**Sparsity analysis.** To test whether connectivity dimensionality reflects sparsity rather than geometric structure, we defined approximate sparsity (since principal component regression produces dense, all-nonzero matrices) as the proportion of weights falling within thresholds near zero (±0.0001, ±0.00005, ±0.00001), then examined its correlation with connectivity dimensionality across regions.

### Activity flow modeling

We implemented activity flow modeling^21,22^ to generate task-evoked activity via network flow. For each subject and target region, source regions were identified from the parcel-level FC graph (as in ’FC estimation’). Source region activity estimates (task conditions × source vertices) were projected onto each target region via matrix multiplication with the dense FC graph (source vertices × target vertices) in each session: each target vertex’s predicted activity is the sum, over all n connected source vertices, of source activity weighted by source-target connectivity strength (i.e., connectivity-specified activity flow from each source vertex). Model evaluation used the Pearson correlation between the flow-generated and empirically observed task condition activity matrices per target region, averaged over task conditions and sessions.

From model-generated activity estimates, we derived predicted RSMs and representational dimensionality, evaluated against observed values using cosine similarity (RSMs) and Pearson correlation (dimensionality) per region. Prediction residuals (observed minus predicted) were examined to identify regions where additional factors beyond connectivity-based flow—e.g., local circuit dynamics, neuromodulation, attention—may contribute.

Significance was estimated by projecting source activity over 100 iterations of randomly shuffled connectivity per subject, generating 1000 group-level bootstrapped means ^63^. A more stringent test constrained shuffling to the source side only—randomly permuting connectivity values among vertices within each source region—preserving each source’s spatial distribution properties and leaving target-vertex connectivity properties unaltered. Significance against this null demonstrates that specific source-to-target connectivity patterns, rather than general properties like target vertex degree distribution, drive predictive accuracy.

### Double cross-validated RSMs for cortical dimensionality pattern comparison

To test whether model-predicted dimensionality patterns replicate the compression-then-expansion pattern along the sensorimotor FC gradient, we developed double-cross-validated RSMs, cross-validating between observed and model-predicted estimates in addition to between sessions. Specifically, we computed cosine similarity between observed activity for condition A (session 1) and model-predicted activity for condition B (session 2), yielding a task conditions × task conditions RSM, and repeated this for the reverse combination (predicted session 1 vs. observed session 2). Representational dimensionality was computed from each and averaged within each region, providing a stringent test that reduces noise and session-specific artifacts.

### RSM-based transformation index

We developed an RSM-based transformation index to quantify how much model-generated target representations differ from observed source representations. For each target region we computed three cosine distances (1 − cosine similarity), averaged across source-target pairs: (a) dtransform, between observed source and observed target RSMs; (b) d transform, between observed source and predicted target RSMs; and (c) dpred, between predicted and observed target RSMs (a reference measure of prediction error).

### Cross-task representational similarity

To isolate cross-task from within-task similarity, we performed a one-sample t-test on each RSM cell across subjects (FDR-corrected), yielding significance-survived RSMs of reliable task-condition combinations. From these, we computed average positive similarity values among survived combinations (restricted to positive values, since negative cosine similarities are difficult to interpret), providing a measure of cross-task representational relationships.

To assess this relationship more generally, we simulated activity propagation across connectivity matrices with varying dimensionality. Source and target region sizes were sampled from uniform distributions of (100,150) and (50,100) vertices, respectively. For each source region, 100 random task activity patterns (standard normal distribution) were propagated through low-rank connectivity matrices (also standard normal) at rank factors of [0.01, 0.025, 0.05, 0.1, 0.2, 0.4, 0.6, 0.8, 1.0] × target region size (10 source-target pairs per rank factor). Cross-task similarity was computed as the cosine similarity between resulting target task activity patterns, averaging positive values in the upper-triangular similarity matrix.

### Representational similarity analysis in multiple tasks with shared features

We identified five tasks (44 task conditions) with especially interpretable shared features to test whether conjunctive feature coding relates to representational expansion. We constructed ground-truth similarity matrices for sensory, cognitive, and motor features, and their multiplicative conjunction.

Sensory and motor features were coded from stimuli and expected responses in each task condition (Supplementary Data Table 1). Stimuli were categorized hierarchically (higher-order features: text, images, videos, spatial arrays; specific elements: scenes, faces, objects), yielding a graded stimulus RSM (1 = same stimulus type 1, e.g. images; 2 = same stimulus type 2, e.g. objects; 3 = identical stimuli reused across conditions). Given practiced task accuracy (>85%) prior to scanning, we constructed a binary motor similarity matrix from expected button-press responses (1 = same response, 0 = different).

Cognitive (task demand) similarity was computed by assigning demand levels to each task’s cognitive manipulation—most tasks had three levels (easy/medium/hard, coded 1–3), with Stroop coded 1 (congruent) or 3 (incongruent)—and computing pairwise similarity as:

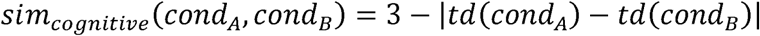

where *td* represents task demand, yielding values between one and three.

All three matrices were normalized to [0,1] and multiplied to produce the conjunction term. In each region and subject, the observed 44×44 RSM (averaged across subjects for visualization) was fit to the four ground-truth matrices using ridge regression with 5-fold cross-validation.

### Human Connectome Project: neural and behavioral data analysis

We analyzed 352 unrelated, low-motion subjects from the HCP ’1200 Subjects’ release, computing representational dimensionality from HCP task data^27,28^ using procedures analogous to our primary dataset. Following the same preprocessing pipeline, we obtained activity estimates for 24 task conditions across seven tasks via GLM-based multiple regression, with separate regressors per condition across two identical runs differing only in phase-encoding direction. Cross-validated RSMs (cosine similarity between runs) yielded dimensionality metrics, and a compression ratio quantified relative compression in association versus sensory/motor regions:

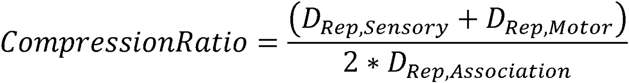

where D_Rep_ represents the average representational dimensionality for each region type.

For behavioral analysis, we used a general fluid intelligence composite score^64^, combining the z-scored fluid cognition composite (Dimensional Change Card Sort, Flanker, Picture Sequence Memory, List Sorting, Pattern Comparison^65,66^) with z-scored Penn Matrix Reasoning Test performance, reflecting broad cognitive ability across working memory, cognitive control, and reasoning^47^.

## Data Availability

The activation maps and functional parcellations from the original study are available from http://www.diedrichsenlab.org/imaging/mdtb.htm. The raw behavioral and imaging data for the cerebellum are also available on the data sharing repository https://openneuro.org/. Example derivatives (representational and connectivity dimensionality values of all subjects) can be accessed at https://github.com/LakshmanChakravarthy/multitask_generalization/tree/master/example_deriva tives.

## Code Availability

The codebase can be accessed at https://github.com/LakshmanChakravarthy/multitask_generalization.

## Extended Data

**Extended Data Figure 1:**
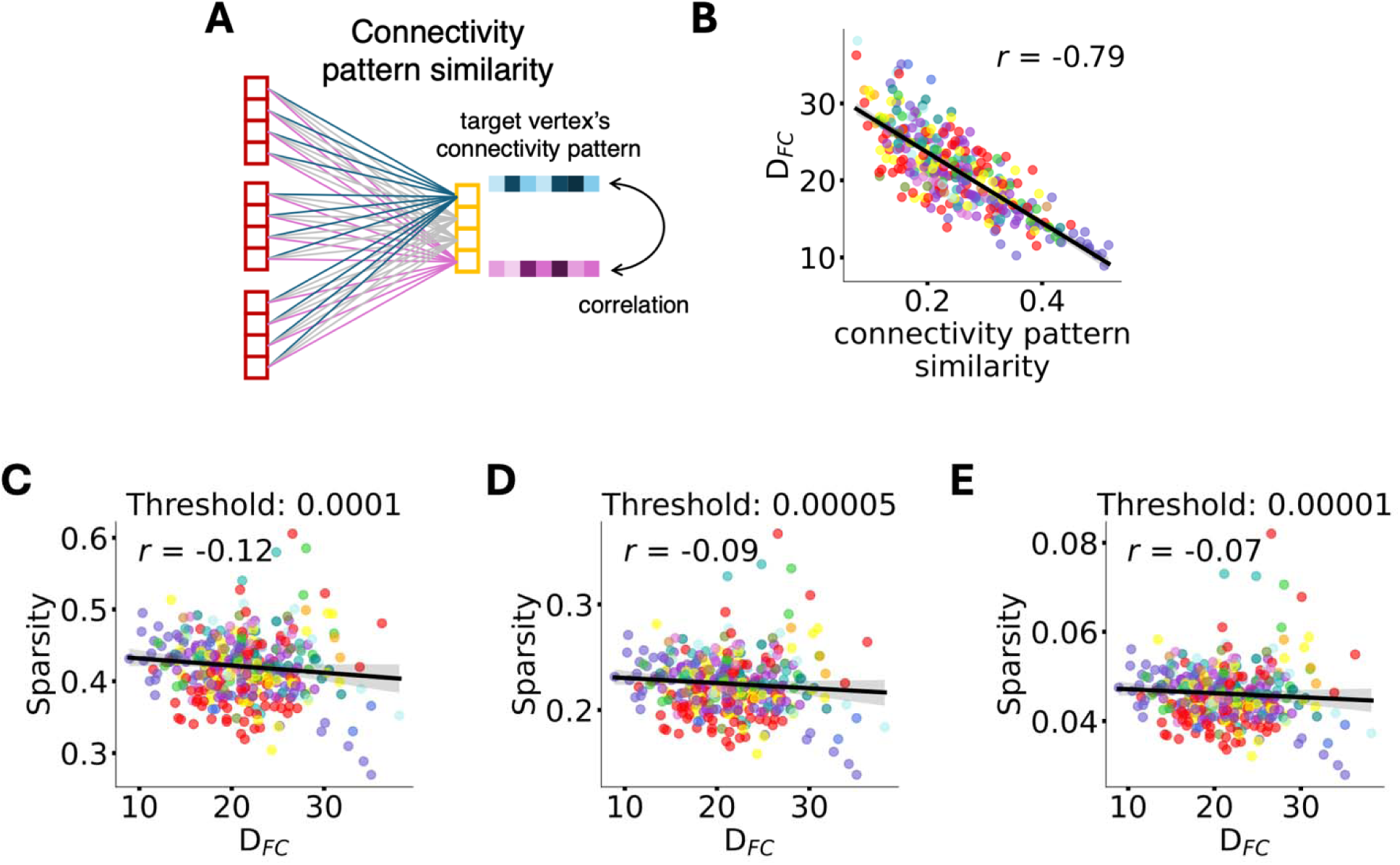
Connectivity dimensionality reflects similarity of connectivity patterns, not sparsity. A) Schematic of connectivity pattern similarity. Each target vertex (yellow and purple boxes on right) receives weighted inputs from all source vertices (red boxes on left), forming a connectivity pattern (vector of connection weights). Connectivity pattern similarity is measured by computing the pairwise Pearson correlation between all target vertices’ connectivity patterns within each region, then averaging these correlations. **B) Connectivity dimensionality is strongly determined by connectivity pattern similarity.** Correlation between connectivity pattern similarity (mean pairwise correlation among target vertices’ connectivity patterns) and connectivity dimensionality (DFC) across all 360 regions (r = −0.79, p < 0.001, spatial-autocorrelation preserving permutation test). Each dot represents one region, colored by network. When target vertices have similar connectivity patterns (high correlation), dimensionality is low. When target vertices have diverse connectivity patterns (low correlation), dimensionality is high. This demonstrates that connectivity dimensionality captures the geometric structure of how regions connect, not merely the number or strength of connections. **C-E) Connection sparsity does not correlate with connectivity dimensionality.** Principal component regression produces dense connectivity matrices (all weights non-zero). To test whether approximately sparse connectivity (many near-zero weights) explains dimensionality, we computed sparsity as the proportion of connection weights falling within small thresholds around zero: ±0.0001 (panel C), ±0.00005 (panel D), and ±0.00001 (panel E). Across all three thresholds, sparsity showed no significant relationship with connectivity dimensionality (all p > 0.05). This confirms that connectivity dimensionality reflects the similarity structure among connectivity patterns rather than connection density.

**Extended Data Figure 2:**
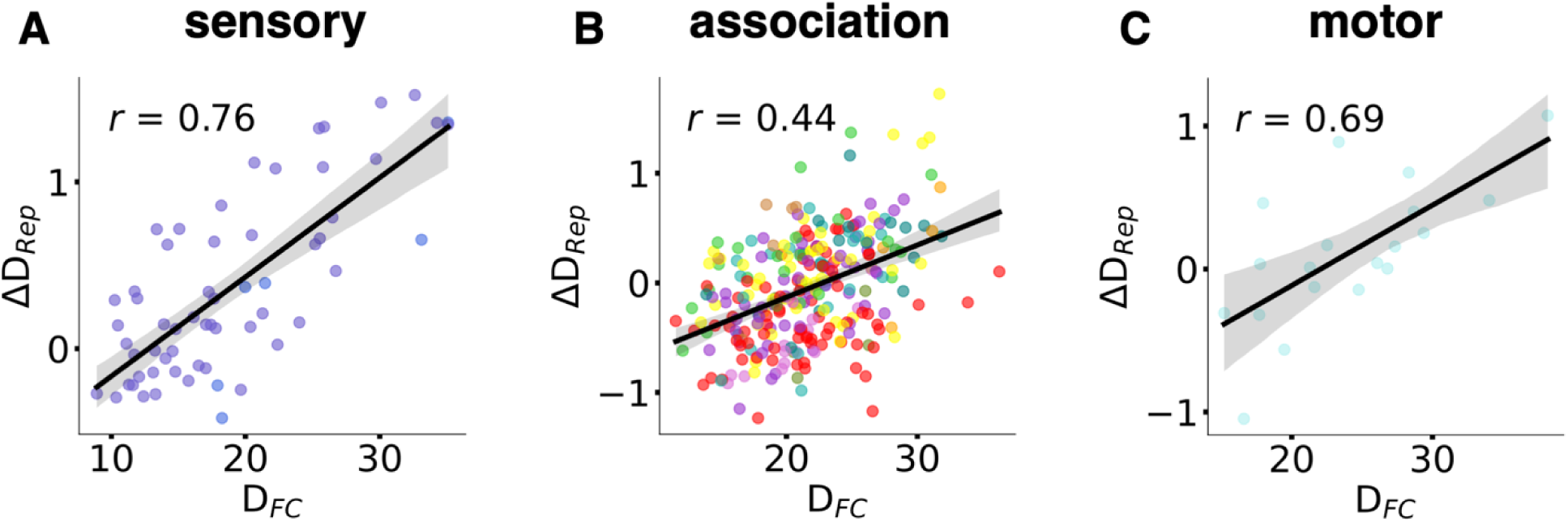
Connectivity dimensionality predicts compression/expansion within sensory, association, and motor systems. The relationship between connectivity dimensionality (DFC) and representational compression/expansion (ΔDRep) holds consistently across all three major cortical divisions. **A) Sensory regions** (visual networks VIS1 and VIS2; n=59 regions): r = 0.76, p < 0.001. **B) Association regions** (default-mode, frontoparietal, dorsal attention, language, cingulo-opercular, auditory, and somatomotor-parietal networks; n=250 regions): r = 0.44, p < 0.001. **C) Motor regions** (somatomotor-frontal network SMN-Fr; n=51 regions): r = 0.69, p = 0.0012. Each dot represents one region, colored by network identity within each system. Gray shading shows 95% confidence interval of the linear fit. These within-system correlations demonstrate that connectivity geometry predicts representational transformations as a general principle operating across functionally distinct cortical systems, not merely as a whole-brain statistical artifact.

**Extended Data Figure 3:**
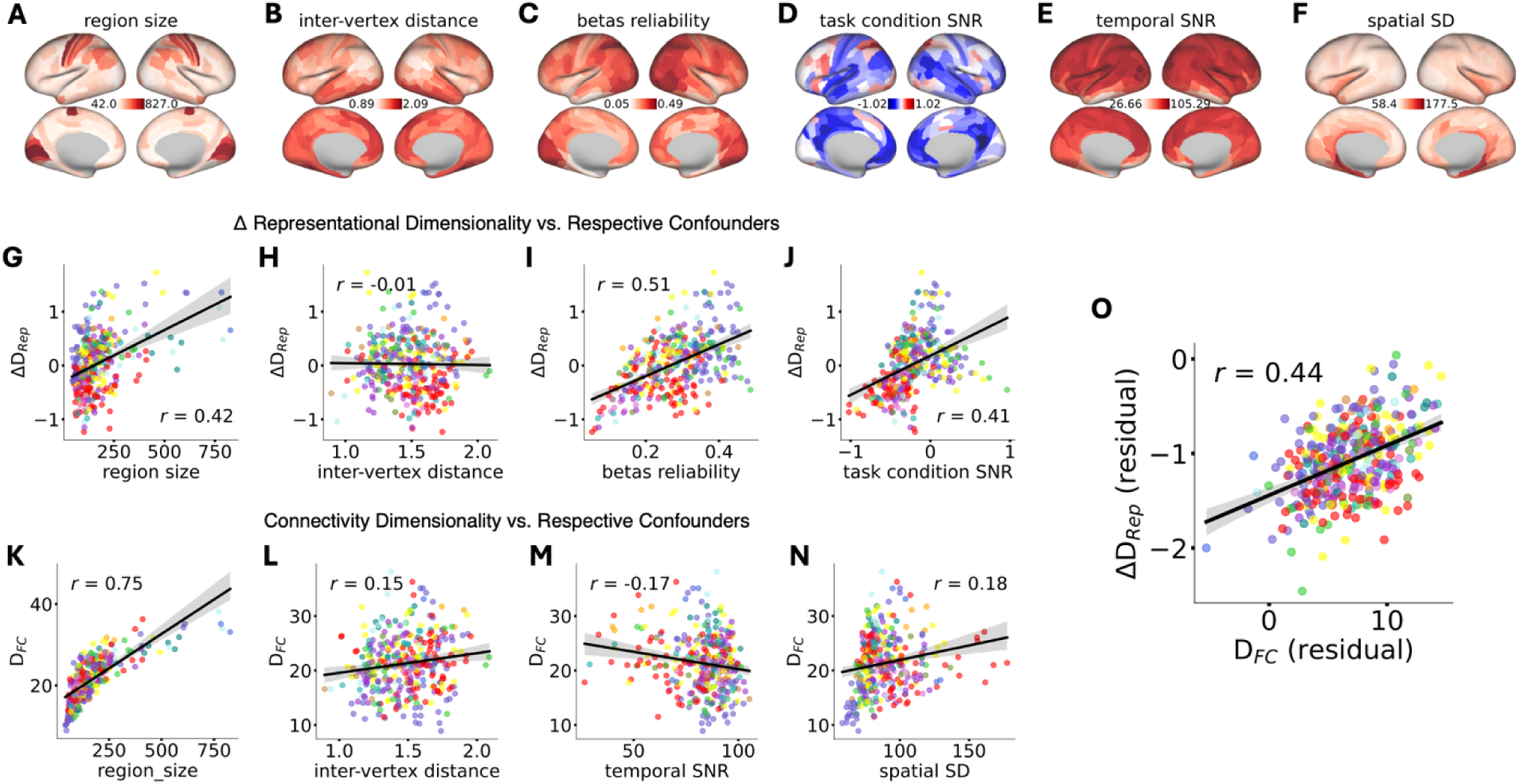
Relationship between connectivity dimensionality and representational compression/expansion remains significant after controlling for potential confounds. A-F) Spatial distribution of potential confounding variables across cortex. Brain maps showing six region-specific properties varying across cortex that could potentially confound connectivity dimensionality, representational dimensionality change, or both. **A) Region size** (vertex count). **B) Inter-vertex distance** (mean geodesic distance on cortical surface). **C) Betas reliability** (mean task condition reliability, estimated by taking the mean of the RSM diagonal). **D) Task Condition Signal-to-Noise Ratio, tcSNR** (mean/SD of task condition betas at each vertex, averaged across vertices in each region). **E) Temporal Signal-to-Noise Ratio, tSNR** (mean/SD of rest time series at each vertex, averaged across vertices in each region). **F) Spatial variance** (standard deviation of rest time series across vertices at each timepoint, averaged across time). **G-J) Individual confound relationships with compression/expansion.** Each panel shows correlation between one confound and ΔD_Rep_ across all 360 regions. Region size (r = 0.42), betas reliability (r = 0.51), and task condition SNR (r = 0.41) show significant relationships; inter-vertex distance shows no relationship (r = −0.01). **K-N) Individual confound relationships with connectivity dimensionality.** Each panel shows correlation between one confound and D_FC_ across all 360 regions. Region size shows the strongest relationship with D_FC_ (r = 0.75), while inter-vertex distance (r = 0.15), temporal SNR (r = −0.17), and spatial SD (r = 0.18) show smaller but significant relationships. These correlations demonstrate that several region-specific properties do vary with compression/expansion and connectivity dimensionality, necessitating control analyses. **O) Central result remains significant after controlling for confounds.** To test whether connectivity dimensionality independently predicts compression/expansion, we regressed the respective confounds from ΔD_Rep_ and D_FC_ and correlated the residuals, and the resultant correlation is still significant (r = 0.4357, p<0.001). These results demonstrate that connectivity dimensionality predicts representational transformations independent of region size, inter-vertex distance, beta reliability, task condition SNR, temporal SNR, and spatial variance.

**Extended Data Figure 4:**
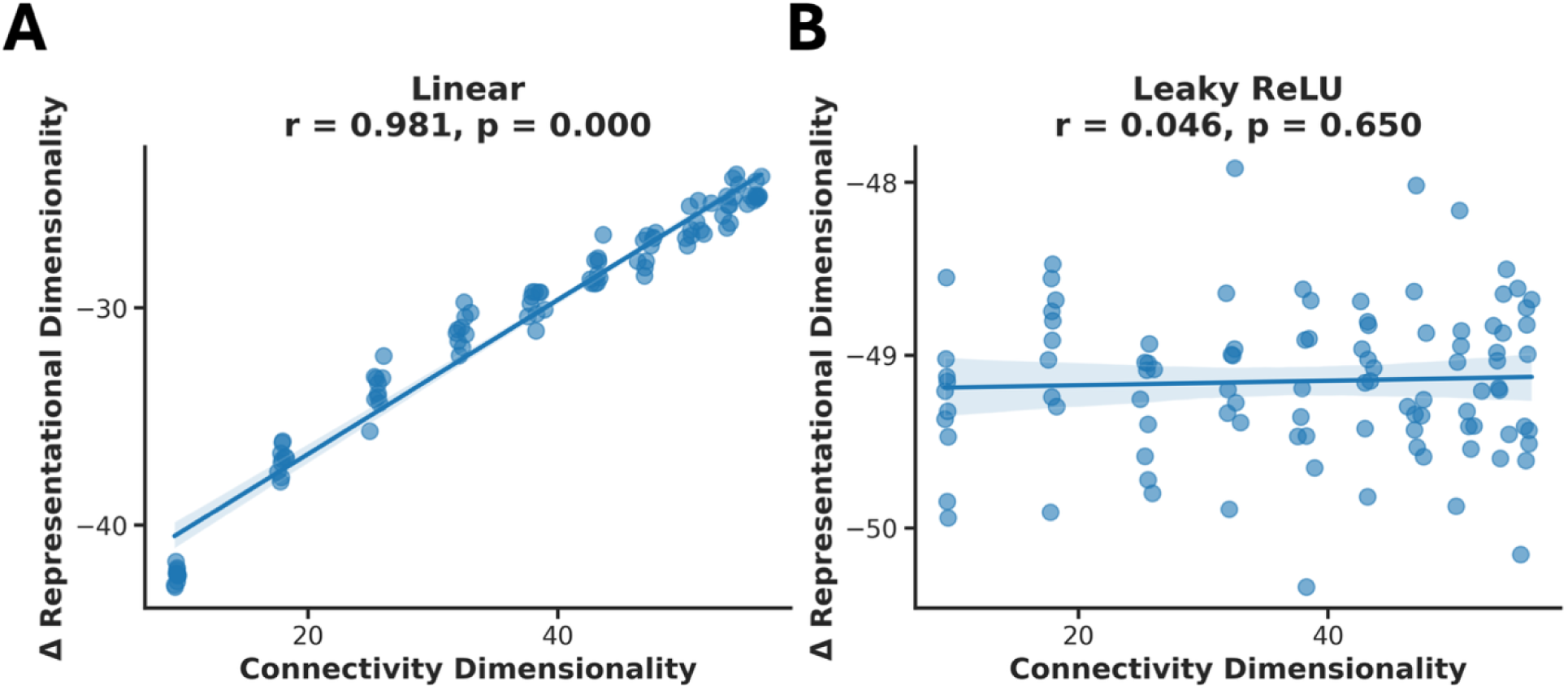
Non-linear local computations can decouple connectivity dimensionality from representational dimensionality change. To test whether this relationship is mathematically inevitable, we simulated source and target regions (100 vertices each) with varying connectivity dimensionality. Source activity patterns for 100 tasks were generated from a standard normal distribution and projected through connectivity matrices with dimensionality controlled by rank (participation ratio ranging from 10–100). **A) Linear projection preserves the relationship.** When source activity is linearly projected through the connectivity matrix and representational dimensionality is computed directly from target activity, the relationship between connectivity dimensionality and change in representational dimensionality (target minus source) is extremely strong (r = 0.981, p < 0.001). Each dot represents one simulated source-target pair (10 pairs per dimensionality level, across 9 levels). Lower connectivity dimensionality produces compression (negative Δ representational dimensionality); higher dimensionality produces less compression or expansion, demonstrating that connectivity dimensionality strongly constrains representational dimensionality under purely linear transformations. **B) Non-linear local computations eliminate the relationship.** To simulate nonlinear transformation at the target region, connectivity-projected activity additionally underwent a leaky ReLU non-linearity (alpha = 0.01) with region-specific and task-dependent bias terms before representational dimensionality was computed: target vertex activity, where source task activity 0,1 , is leaky ReLU with = 0.01, is the trainable parameter, but in this case are connectivity weights with set dimensionality, bias pattern that simulates idiosyncratic local computations in the target region, where each vertex has a baseline bias drawn from a Gaussian distribution with a large mean for demonstration, 400,50 , and task-specific biases with smaller variance on top, 0,25 . When simulated this way, the relationship between compression and connectivity dimensionality disappears (r = 0.046, p = 0.650). This exaggerated scenario demonstrates that strong non-linear local operations can override connectivity-based constraints on representational dimensionality, and that such relationship is not mathematically inevitable. The empirical observation that connectivity dimensionality robustly predicts representational transformations across cortex (Fig. 2E, r = 0.44) therefore suggests that while local computations certainly contribute, connectivity geometry provides systematic constraints that shape representational transformations throughout the brain. Note that this linear projection assumption applies to the activity flow model, not to the primary empirical result; it does not imply that neuronal transformations are themselves linear, but rather that linear projection approximates their net effect at the fMRI spatial scale.

**Extended Data Figure 5:**
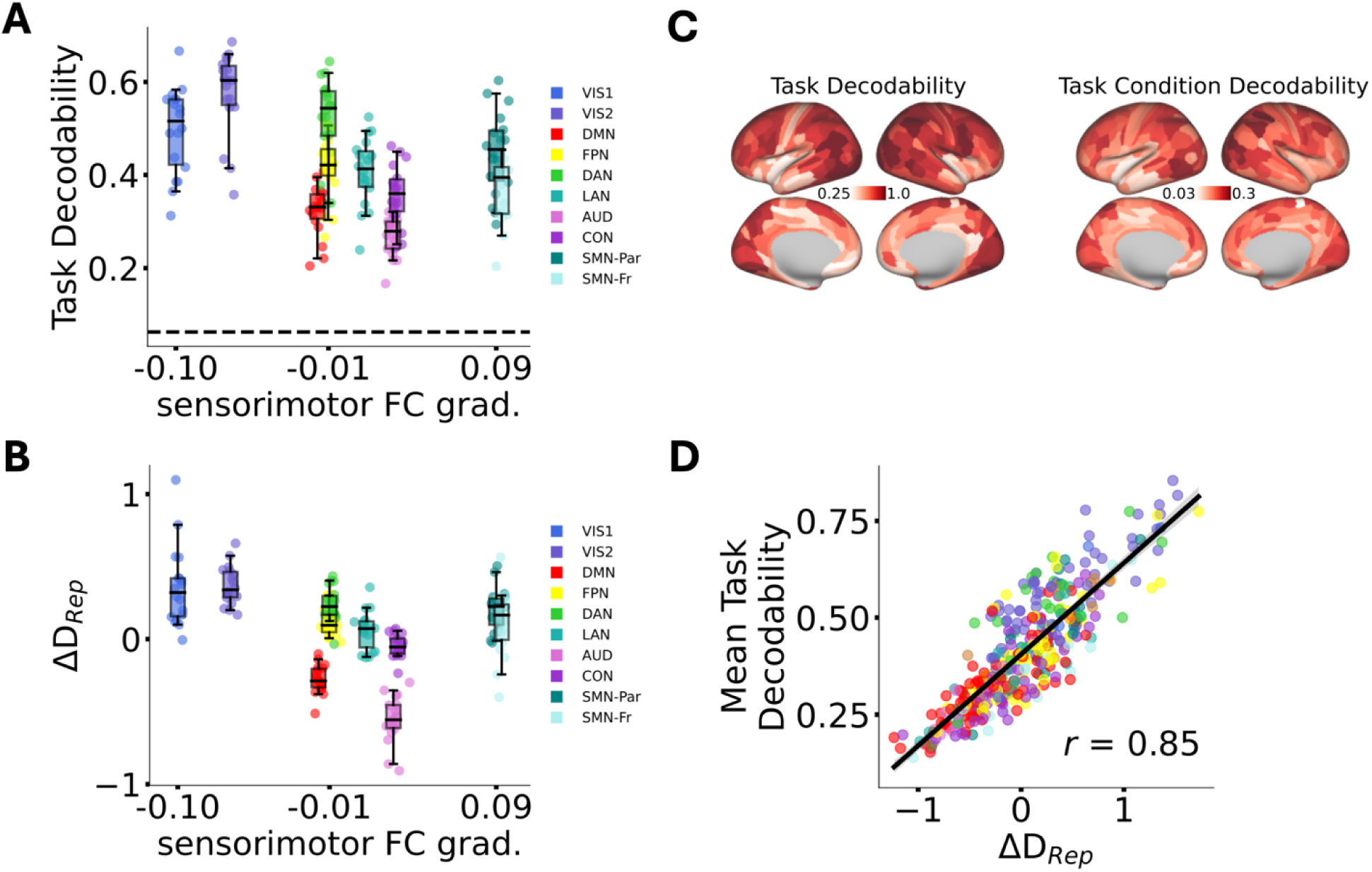
Representational compression corresponds to reduced task decodability across cortical networks. *Motivation:* BOLD signal in association cortex, where mixed-selective neurons are spatially intermingled, may poorly reflect underlying representational diversity, consistent with prior reports of difficulty decoding task information from prefrontal BOLD^47^. **A) Task decodability varies across cortical networks.** Decodability of 16 individual tasks from representational similarity matrices (2-fold minimum distance classification based on cosine distance from task-averaged RSMs; chance = 1/16 = 0.0625, dashed line). Each dot represents one region; box plots show distribution across regions within each network arranged along the sensory-motor gradient. Visual networks (VIS1, VIS2) and motor regions (SMN-Fr) show highest task decodability. Default-mode (DMN) and auditory (AUD) networks show lowest decodability, falling near or below the median of other networks. **B) Compression/expansion pattern mirrors decodability.** Representational compression/expansion (ΔD_Rep_) plotted by network along the sensory-motor gradient. Default-mode and auditory networks show strongest compression (most negative ΔD_Rep_), while visual and motor networks show expansion or minimal compression. This pattern closely parallels the decodability results in panel A, demonstrating that compression reflects genuine loss of task discriminability. **C) Task decodability is robust throughout cortex.** Brain maps showing decodability of 16 tasks (left; range 0.25-1.0) and 96 task conditions (right; range 0.03-0.3). Despite the compression observed in association cortex, task decodability remains well above chance (0.0625) even in default-mode regions, indicating that BOLD signals in association cortex meaningfully reflect task information rather than averaging over mixed selectivity to the point of indiscriminability. **D) Mean task decodability strongly correlates with compression/expansion.** Correlation between mean task decodability and ΔDRep across all 360 regions (r = 0.85, p < 0.001). Each dot represents one region, colored by network. Regions showing expansion (positive ΔD_Rep_) have higher task decodability; regions showing compression have lower decodability.

**Extended Data Figure 6:**
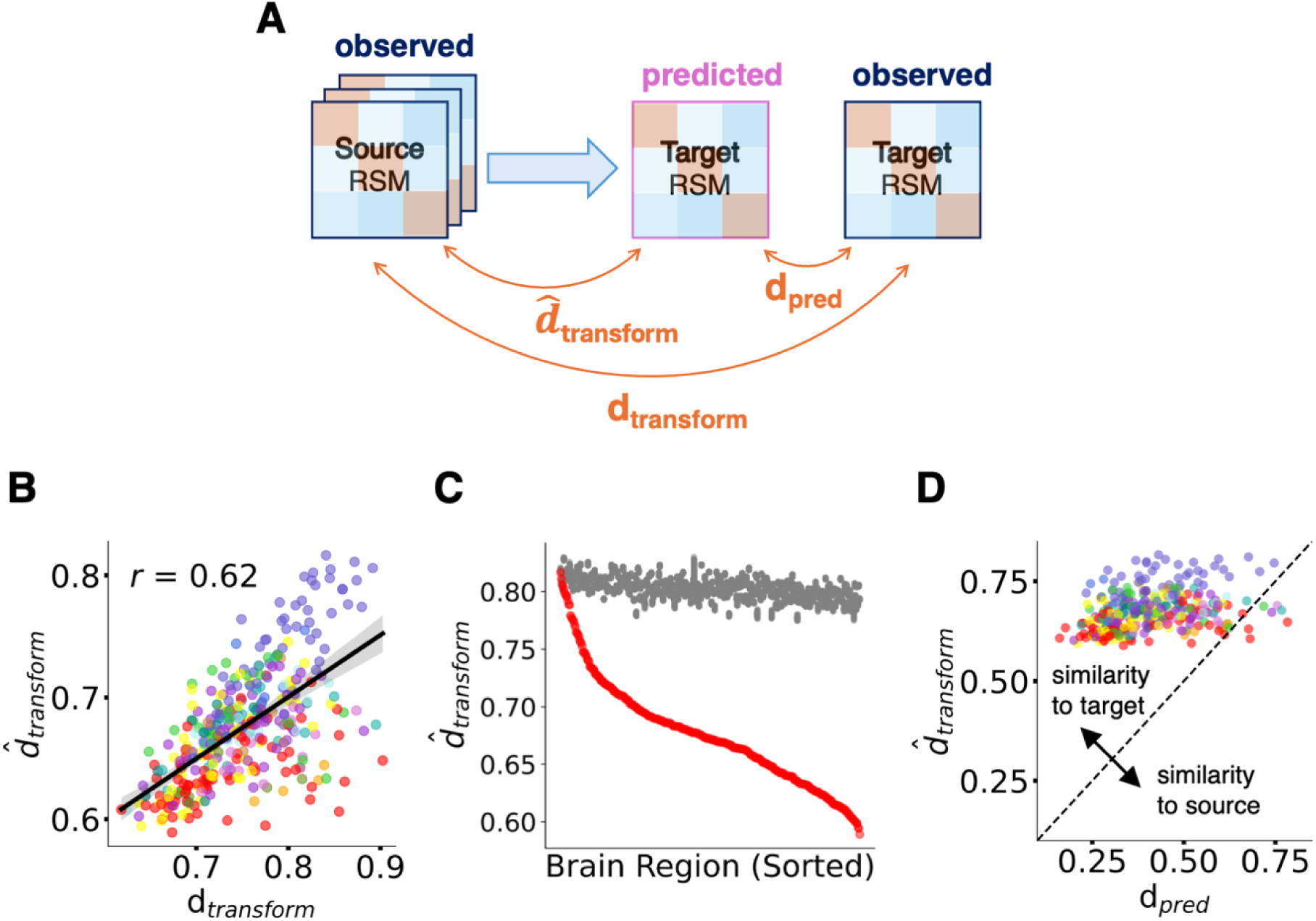
Activity flow over connectivity generates genuine representational transformations, not simple transfer. A) Measuring transformation distance. To test whether activity flow implements true representational transformations rather than simply transferring source representations to targets, we computed three distances using cosine distance (1 - cosine similarity) between RSMs. d **_transform_**: distance between observed source RSM and predicted target RSM, quantifying how much the model transforms representations. **d_transform_**: distance between observed source RSM and observed target RSM, quantifying the actual transformation. **d_pred_**: distance between predicted target RSM and observed target RSM, quantifying prediction error. **B) Predicted transformation distance correlates with observed transformation distance.** Correlation between d _transform_ (model-predicted source-to-target transformation) and d_transform_ (observed source-to-target transformation) across all 360 regions (r = 0.62, p < 0.001, spatial-autocorrelation preserving permutation test). Each dot represents one region, colored by network. The model accurately captures the magnitude of representational transformations across cortex. **C) Connectivity-based transformations differ significantly from null models.** Predicted transformation distances (d _transform_, red) plotted for each region sorted by magnitude, compared against null distributions generated by shuffled connectivity (gray; 1000 iterations). All 360 regions show significantly lower transformation distances with intact connectivity than with shuffled connectivity (all p < 0.001), demonstrating that the specific structure of empirical connectivity patterns drives transformations. **D) Predictions resemble targets more than sources in most regions.** Each dot represents one region, plotting prediction distance (d_pred_, similarity to target) against predicted transformation distance (d _transform_, similarity to source). Points below the diagonal (339/360 regions, pFDR < 0.05) indicate the predicted RSM is closer to the observed target than to the observed source, demonstrating genuine transformation rather than simple transfer. This holds for both compression zones (association cortex) and expansion zones (motor cortex), showing that connectivity geometry shapes transformations bidirectionally.

**Extended Data Figure 7:**
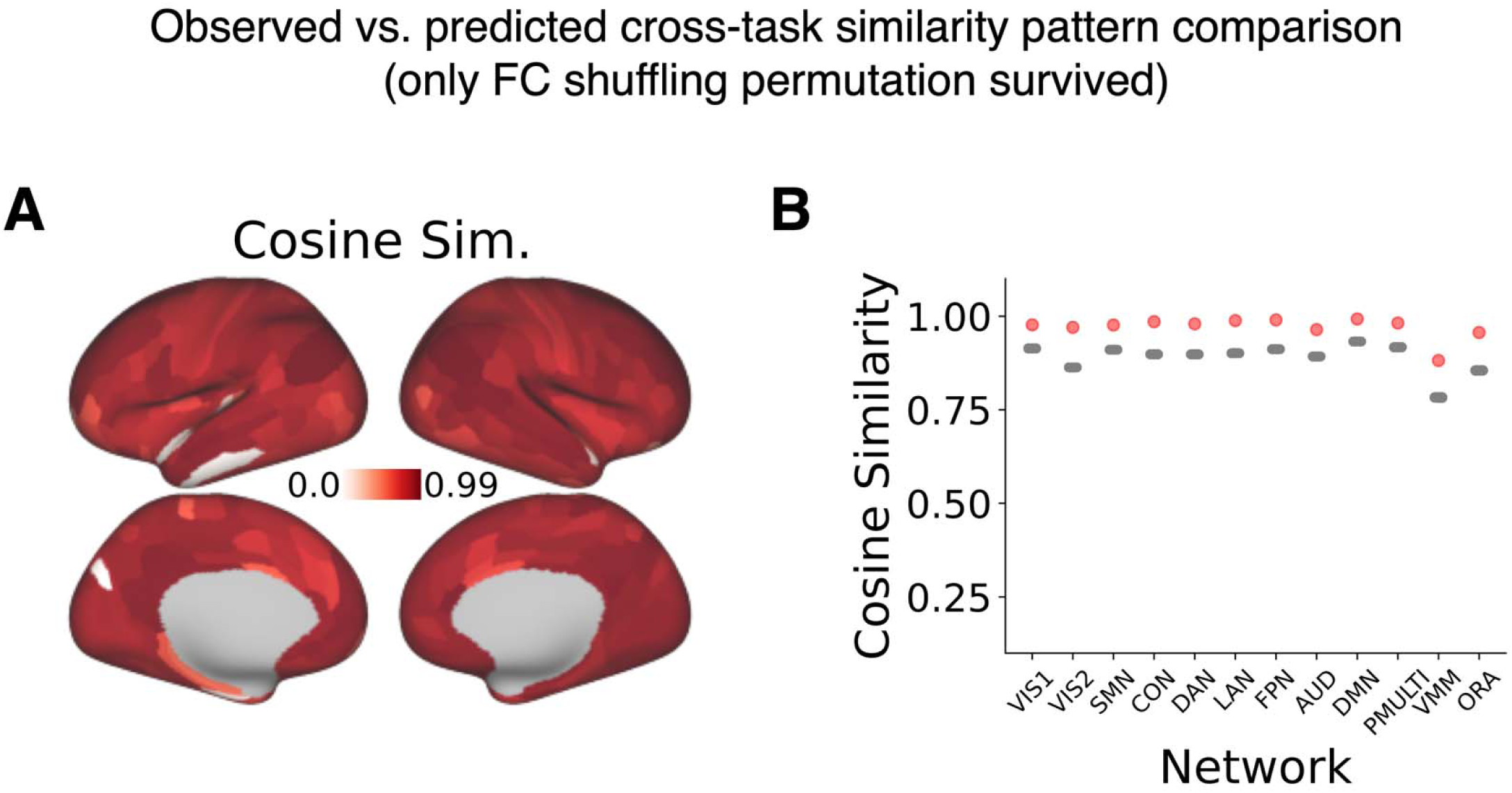
Cross-task similarity predictions at the level of individual regions and large-scale brain networks. (A) Cosine similarity between vectorized cross-task similarity values from observed (empirically-estimated) and from connectivity-predicted estimates (range 0.13–0.99; p_connperm_ < 0.05 in 355/360 regions). Regions with no values did not survive FC shuffling permutation test. (B) Similar comparison of the network-level cross-task similarity patterns. Grey dots indicate null distributions derived from FC shuffling. Network abbreviations from ^27^: VIS1: primary visual network, VIS2: secondary (higher-order) visual network, SMN: somato-motor network, CON: cingulo-opercular network, DAN: dorsal attention network, LAN: language network, FPN: fronto-parietal network, AUD: auditory network, DMN: default-mode network, PMULTI: posterior multimodal network, VMM: ventral multimodal network, ORA: orbito-affective network.

**Extended Data Figure 8:**
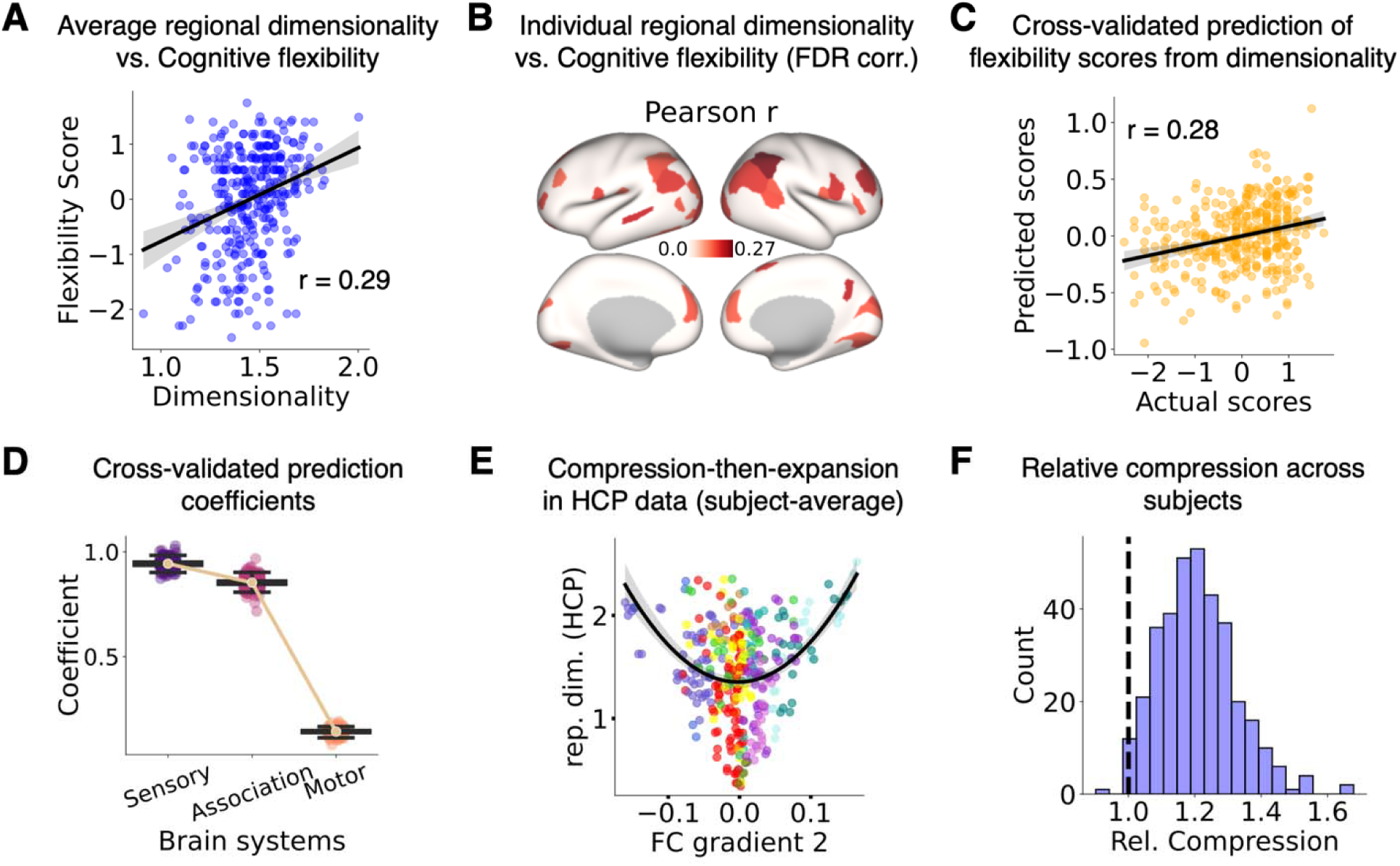
Representational dimensionality is predictive of behavioral measures of cognitive flexibility. A) Average representational dimensionality across regions was positively correlated with cognitive flexibility scores across subjects. B) Similar relationship in individual regions (following multiple comparison corrections) was reflected in positive correlations in different regions. C) Dimensionality values averaged across sensory, association and motor regions were predictive of flexibility scores in a cross-validated (leave one subject out) manner. D) Coefficients from the cross-validation approach show higher values for sensory and association regions’ dimensionality values compared to motor regions in explaining the flexibility scores. E) The representational dimensionality values in the new dataset also follow compression-then-expansion pattern (average values across subjects). F) Relative compression in association regions compared to sensory and motor regions was reflected in almost all subjects, measured using compression ratio. These findings address an apparent paradox: although association regions show representational compression on average (Fig. 2), individual differences in absolute dimensionality within this organization—particularly in sensory and association regions—predict fluid intelligence, suggesting the compression-then-expansion pattern reflects canonical task organization while absolute dimensionality reflects individual capacity for flexible cognition.

**Extended Data Table 1:**
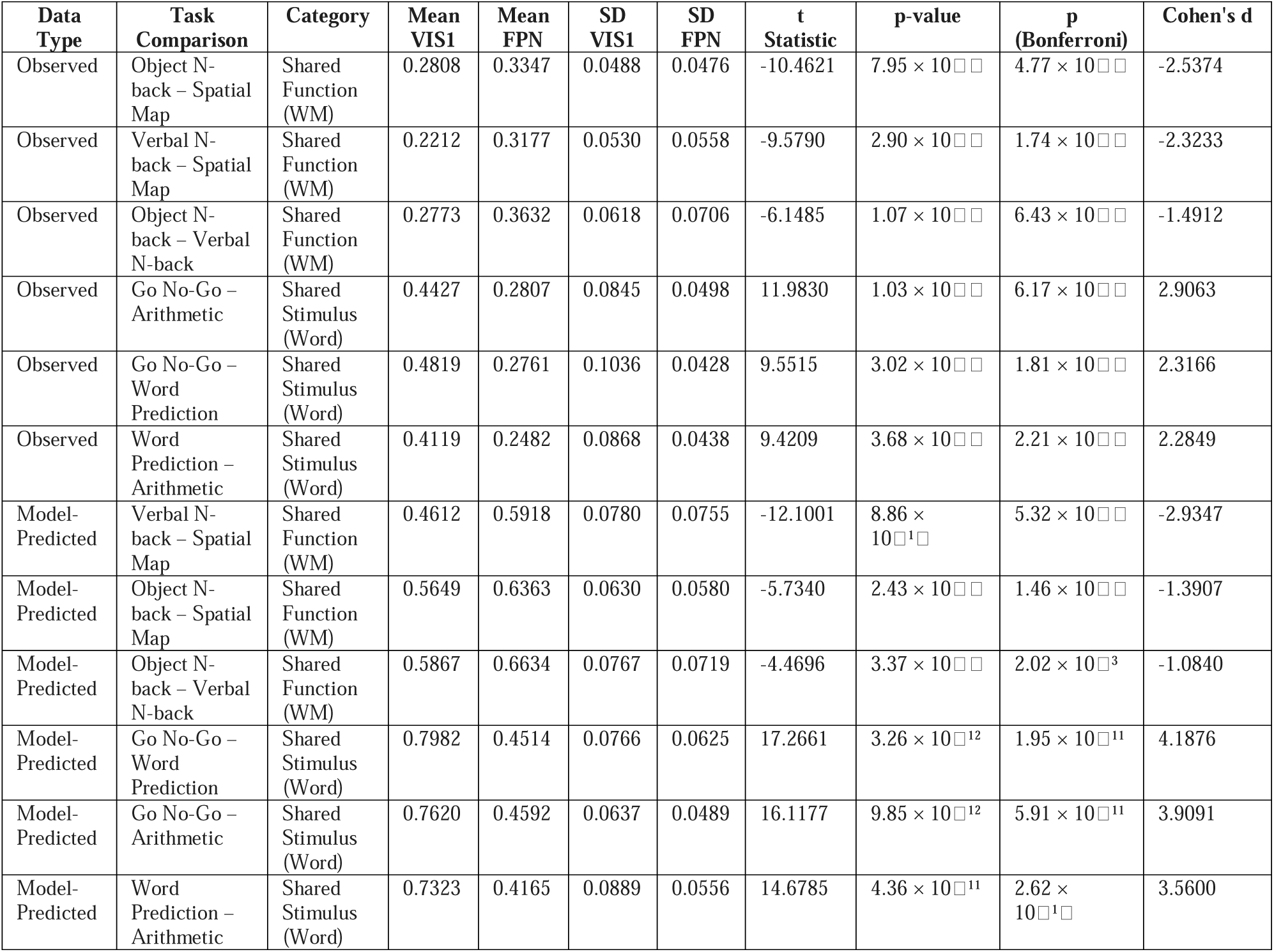
Cross-task similarity comparison between tasks and networks.

**Extended Data Table 2:**
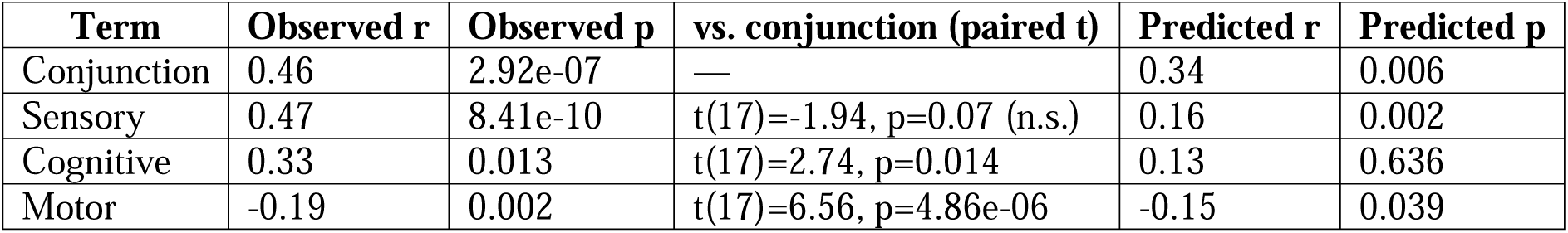
Correlations between representational expansion and RSA template term coefficients, observed and predicted. Anatomical specificity of the four RSA template terms: 9/10 highest sensory coefficients in sensory regions, 7/10 highest cognitive coefficients in association regions, 7/10 highest motor coefficients in motor regions (Fig. 5B). Conjunction-term coefficients correlated with representational expansion (ΔD_Rep_) more strongly than cognitive or motor terms alone, and comparably to the sensory term.

## Supplementary Information

**Supplementary Data Table 1:**
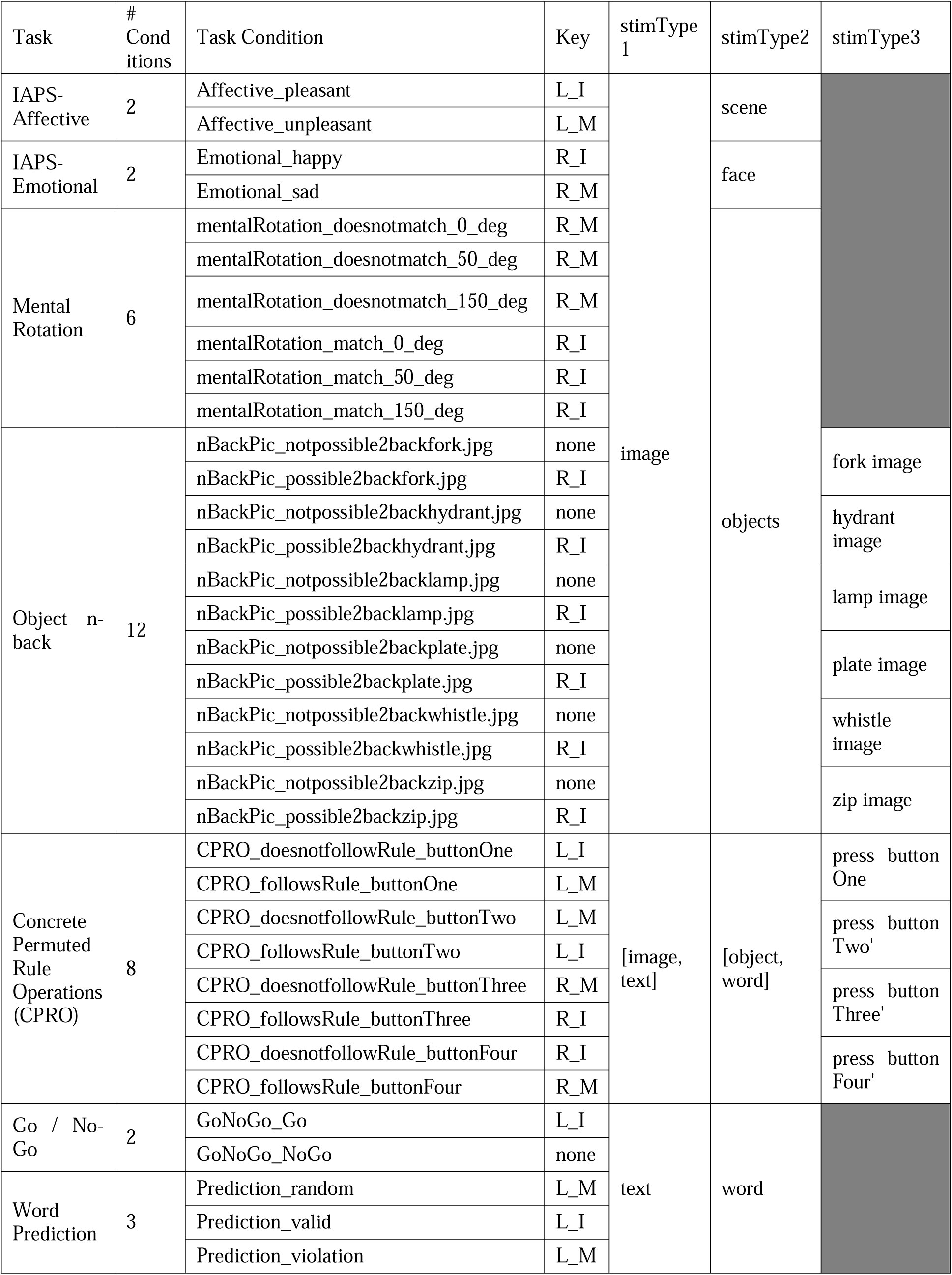
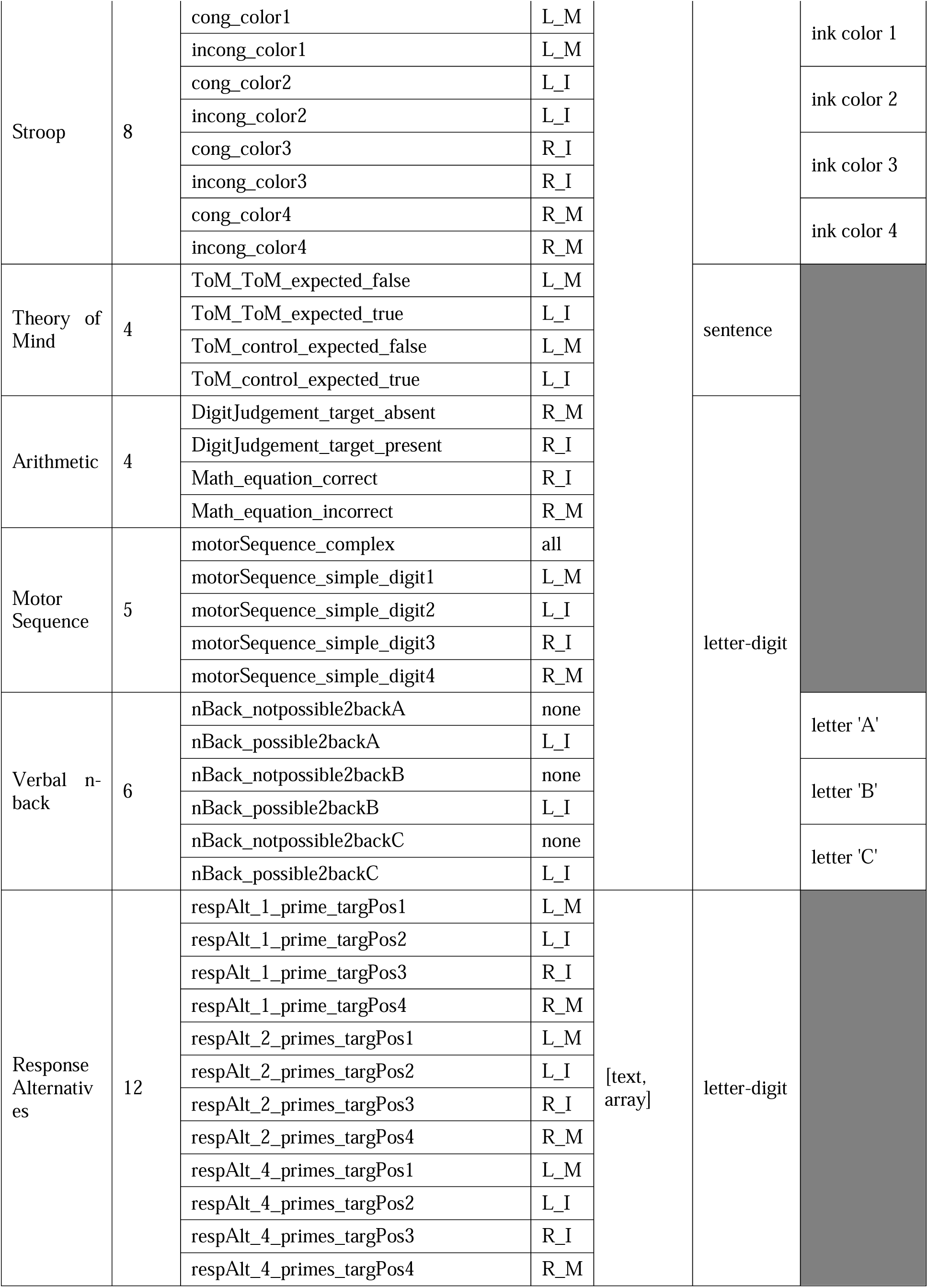
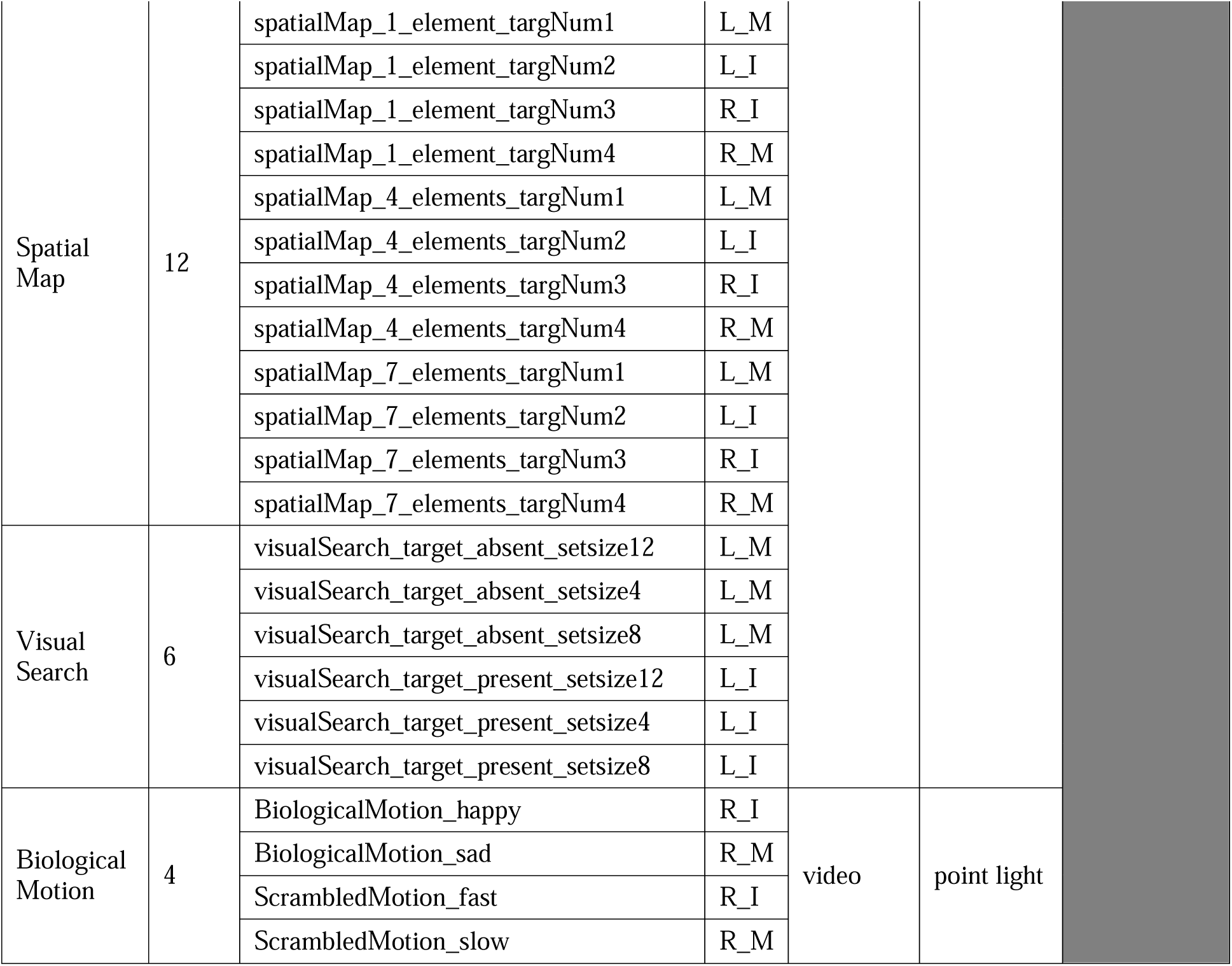
Task condition split of 16 active, visual tasks in Multi-domain Task Battery Dataset. For more information on task description, see the supplementary information of the original paper ^48^. Response key index: L_I: left index finger button, L_M: left middle finger button, R_I: right index finger button, R_M: right middle finger button, all: response using all four buttons was required, none: no response required.

