## Supplementary material for "Network geometry shapes the transformation of task representations across human cortex": Inventory of Supporting Information

**Manuscript #:** NN-A91756B

Please complete each of the Inventory Tables below to outline your Extended Data and Supplementary Information items.

There are four sections:

- *Extended Data*
- *Supplementary Information: PDF Files*
- *Supplementary Information: Additional Files*
- *Source Data*

Each section includes specific instructions. Please complete these tables as fully as possible. We ask that you avoid using spaces in your file names, and instead use underscores, i.e.: Smith_ED_Fig1.jpg not Smith ED Fig1.jpg

Please note that titles and descriptive captions will only be lightly edited, so please ensure that you are satisfied with these prior to submission.

If you have any questions about any of the information contained in this inventory, please contact the journal.

**Corresponding author name(s):** Lakshman Nallan Chakravarthula

1. **Extended Data**

**Complete the Inventory below for all Extended Data Figures and Tables**

- Keep Figure/Table titles to one sentence only
- Upload your files as ‘Figure Files’ in our Manuscript Tracking system
- File names should include the Figure/Table Number. i.e.: *Smith_ED_Fig1.jpg*
- Please be sure to include the file extension in the Filename.
- Note that Extended Data Figures must be submitted as .jpg, .tif, .png, .pdf or .eps files, in RGB colour space, and should be no more than 10MB
- Extended Data Tables may be submitted in .csv, .xlsx or .docx format. Please avoid colours and split or merged cells.
- All Extended Data item legends must be provided in the Inventory below and should not exceed 300 words each *(if possible)*
- Please include Extended Data *ONLY* in this table

| Figure or Table #  Please group Extended Data items by type, in sequential order. Total number of items (Figs. + Tables) must not exceed 10. | Figure/Table title  One sentence only | Filename  Whole original file name including extension. i.e.: Smith_ED_Fig1.jpg | Figure/Table Legend  If you are citing a reference for the first time in these legends, please include all new references in the main text Methods References section, and carry on the numbering from the main References section of the paper. If your paper does not have a Methods section, include all new references at the end of the main Reference list. |
| --- | --- | --- | --- |
| Extended Data Fig. 1 | **Connectivity dimensionality reflects similarity of connectivity patterns, not sparsity.** | ExtendedDataFigure1.png | **A) Schematic of connectivity pattern similarity.** Each target vertex (yellow and purple boxes on right) receives weighted inputs from all source vertices (red boxes on left), forming a connectivity pattern (vector of connection weights). Connectivity pattern similarity is measured by computing the pairwise Pearson correlation between all target vertices' connectivity patterns within each region, then averaging these correlations. **B) Connectivity dimensionality is strongly determined by connectivity pattern similarity.** Correlation between connectivity pattern similarity (mean pairwise correlation among target vertices' connectivity patterns) and connectivity dimensionality (DFC) across all 360 regions (r = -0.79, p < 0.001, spatial-autocorrelation preserving permutation test). Each dot represents one region, colored by network. When target vertices have similar connectivity patterns (high correlation), dimensionality is low. When target vertices have diverse connectivity patterns (low correlation), dimensionality is high. This demonstrates that connectivity dimensionality captures the geometric structure of how regions connect, not merely the number or strength of connections. **C-E) Connection sparsity does not correlate with connectivity dimensionality.** Principal component regression produces dense connectivity matrices (all weights non-zero). To test whether approximately sparse connectivity (many near-zero weights) explains dimensionality, we computed sparsity as the proportion of connection weights falling within small thresholds around zero: ±0.0001 (panel C), ±0.00005 (panel D), and ±0.00001 (panel E). Across all three thresholds, sparsity showed no significant relationship with connectivity dimensionality (all p > 0.05). This confirms that connectivity dimensionality reflects the similarity structure among connectivity patterns rather than connection density. |
| Extended Data Fig. 2 | **Connectivity dimensionality predicts compression/expansion within sensory, association, and motor systems.** | ExtendedDataFigure2.png | The relationship between connectivity dimensionality (DFC) and representational compression/expansion (ΔDRep) holds consistently across all three major cortical divisions. **A) Sensory regions** (visual networks VIS1 and VIS2; n=59 regions): r = 0.76, p < 0.001. **B) Association regions** (default-mode, frontoparietal, dorsal attention, language, cingulo-opercular, auditory, and somatomotor-parietal networks; n=250 regions): r = 0.44, p < 0.001. **C) Motor regions** (somatomotor-frontal network SMN-Fr; n=51 regions): r = 0.69, p = 0.0012. Each dot represents one region, colored by network identity within each system. Gray shading shows 95% confidence interval of the linear fit. These within-system correlations demonstrate that connectivity geometry predicts representational transformations as a general principle operating across functionally distinct cortical systems, not merely as a whole-brain statistical artifact. |
| Extended Data Fig. 3 | **Relationship between connectivity dimensionality and representational compression/expansion remains significant after controlling for potential confounds.** | ExtendedDataFigure3.png | **A-F) Spatial distribution of potential confounding variables across cortex.** Brain maps showing six region-specific properties varying across cortex that could potentially confound connectivity dimensionality, representational dimensionality change, or both. **A) Region size** (vertex count). **B) Inter-vertex distance** (mean geodesic distance on cortical surface). **C) Betas reliability** (mean task condition reliability, estimated by taking the mean of the RSM diagonal). **D) Task Condition Signal-to-Noise Ratio, tcSNR** (mean/SD of task condition betas at each vertex, averaged across vertices in each region). **E) Temporal Signal-to-Noise Ratio, tSNR** (mean/SD of rest time series at each vertex, averaged across vertices in each region). **F) Spatial variance** (standard deviation of rest time series across vertices at each timepoint, averaged across time). **G-J) Individual confound relationships with compression/expansion.** Each panel shows correlation between one confound and ΔD_Rep_ across all 360 regions. Region size (r = 0.42), betas reliability (r = 0.51), and task condition SNR (r = 0.41) show significant relationships; inter-vertex distance shows no relationship (r = -0.01). **K-N) Individual confound relationships with connectivity dimensionality.** Each panel shows correlation between one confound and D_FC_ across all 360 regions. Region size shows the strongest relationship with D_FC_ (r = 0.75), while inter-vertex distance (r = 0.15), temporal SNR (r = -0.17), and spatial SD (r = 0.18) show smaller but significant relationships. These correlations demonstrate that several region-specific properties do vary with compression/expansion and connectivity dimensionality, necessitating control analyses. **O) Central result remains significant after controlling for confounds.** To test whether connectivity dimensionality independently predicts compression/expansion, we regressed the respective confounds from ΔD_Rep_ and D_FC_ and correlated the residuals, and the resultant correlation is still significant (r = 0.4357, p<0.001). These results demonstrate that connectivity dimensionality predicts representational transformations independent of region size, inter-vertex distance, beta reliability, task condition SNR, temporal SNR, and spatial variance. |
| Extended Data Fig. 4 | **Non-linear local computations can decouple connectivity dimensionality from representational dimensionality change.** | ExtendedDataFigure4.png | To test whether this relationship is mathematically inevitable, we simulated source and target regions (100 vertices each) with varying connectivity dimensionality. Source activity patterns for 100 tasks were generated from a standard normal distribution and projected through connectivity matrices with dimensionality controlled by rank (participation ratio ranging from 10–100). **A) Linear projection preserves the relationship.** When source activity is linearly projected through the connectivity matrix and representational dimensionality is computed directly from target activity, the relationship between connectivity dimensionality and change in representational dimensionality (target minus source) is extremely strong (r = 0.981, p < 0.001). Each dot represents one simulated source-target pair (10 pairs per dimensionality level, across 9 levels). Lower connectivity dimensionality produces compression (negative Δ representational dimensionality); higher dimensionality produces less compression or expansion, demonstrating that connectivity dimensionality strongly constrains representational dimensionality under purely linear transformations. **B) Non-linear local computations eliminate the relationship.** To simulate nonlinear transformation at the target region, connectivity-projected activity additionally underwent a leaky ReLU non-linearity (alpha = 0.01) with region-specific and task-dependent bias terms before representational dimensionality was computed: target vertex activity $y=f\left( T\cdot W+b_{region}+b_{task} \right)$, where source task activity $T\sim N\left( 0,1 \right)$, $f$ is leaky ReLU with $alpha$ = 0.01, $W$ is the trainable parameter, but in this case are connectivity weights with set dimensionality, bias pattern that simulates idiosyncratic local computations in the target region, where each vertex has a baseline bias drawn from a Gaussian distribution with a large mean for demonstration, $b_{region}\sim N\left( 400,50 \right)$, and task-specific biases with smaller variance on top, $b_{task}\sim N\left( 0,25 \right)$. When simulated this way, the relationship between compression and connectivity dimensionality disappears (r = 0.046, p = 0.650). This exaggerated scenario demonstrates that strong non-linear local operations can override connectivity-based constraints on representational dimensionality, and that such relationship is not mathematically inevitable. The empirical observation that connectivity dimensionality robustly predicts representational transformations across cortex (Fig. 2E, r = 0.44) therefore suggests that while local computations certainly contribute, connectivity geometry provides systematic constraints that shape representational transformations throughout the brain. Note that this linear projection assumption applies to the activity flow model, not to the primary empirical result; it does not imply that neuronal transformations are themselves linear, but rather that linear projection approximates their net effect at the fMRI spatial scale. |
| Extended Data Fig. 5 | **Representational compression corresponds to reduced task decodability across cortical networks.** | ExtendedDataFigure5.png | Motivation: BOLD signal in association cortex, where mixed-selective neurons are spatially intermingled, may poorly reflect underlying representational diversity, consistent with prior reports of difficulty decoding task information from prefrontal BOLD^47^. **A) Task decodability varies across cortical networks.** Decodability of 16 individual tasks from representational similarity matrices (2-fold minimum distance classification based on cosine distance from task-averaged RSMs; chance = 1/16 = 0.0625, dashed line). Each dot represents one region; box plots show distribution across regions within each network arranged along the sensory-motor gradient. Visual networks (VIS1, VIS2) and motor regions (SMN-Fr) show highest task decodability. Default-mode (DMN) and auditory (AUD) networks show lowest decodability, falling near or below the median of other networks. **B) Compression/expansion pattern mirrors decodability.** Representational compression/expansion (ΔD_Rep_) plotted by network along the sensory-motor gradient. Default-mode and auditory networks show strongest compression (most negative ΔD_Rep_), while visual and motor networks show expansion or minimal compression. This pattern closely parallels the decodability results in panel A, demonstrating that compression reflects genuine loss of task discriminability. **C) Task decodability is robust throughout cortex.** Brain maps showing decodability of 16 tasks (left; range 0.25-1.0) and 96 task conditions (right; range 0.03-0.3). Despite the compression observed in association cortex, task decodability remains well above chance (0.0625) even in default-mode regions, indicating that BOLD signals in association cortex meaningfully reflect task information rather than averaging over mixed selectivity to the point of indiscriminability. **D) Mean task decodability strongly correlates with compression/expansion.** Correlation between mean task decodability and ΔDRep across all 360 regions (r = 0.85, p < 0.001). Each dot represents one region, colored by network. Regions showing expansion (positive ΔD_Rep_) have higher task decodability; regions showing compression have lower decodability. |
| Extended Data Fig. 6 | **Activity flow over connectivity generates genuine representational transformations, not simple transfer.** | ExtendedDataFigure6.png | **A) Measuring transformation distance.** To test whether activity flow implements true representational transformations rather than simply transferring source representations to targets, we computed three distances using cosine distance (1 - cosine similarity) between RSMs. **d̂_transform_**: distance between observed source RSM and predicted target RSM, quantifying how much the model transforms representations. **d_transform_**: distance between observed source RSM and observed target RSM, quantifying the actual transformation. **d_pred_**: distance between predicted target RSM and observed target RSM, quantifying prediction error. **B) Predicted transformation distance correlates with observed transformation distance.** Correlation between d̂_transform_ (model-predicted source-to-target transformation) and d_transform_ (observed source-to-target transformation) across all 360 regions (r = 0.62, p < 0.001, spatial-autocorrelation preserving permutation test). Each dot represents one region, colored by network. The model accurately captures the magnitude of representational transformations across cortex. **C) Connectivity-based transformations differ significantly from null models.** Predicted transformation distances (d̂_transform_, red) plotted for each region sorted by magnitude, compared against null distributions generated by shuffled connectivity (gray; 1000 iterations). All 360 regions show significantly lower transformation distances with intact connectivity than with shuffled connectivity (all p < 0.001), demonstrating that the specific structure of empirical connectivity patterns drives transformations. **D) Predictions resemble targets more than sources in most regions.** Each dot represents one region, plotting prediction distance (d_pred_, similarity to target) against predicted transformation distance (d̂_transform_, similarity to source). Points below the diagonal (339/360 regions, pFDR < 0.05) indicate the predicted RSM is closer to the observed target than to the observed source, demonstrating genuine transformation rather than simple transfer. This holds for both compression zones (association cortex) and expansion zones (motor cortex), showing that connectivity geometry shapes transformations bidirectionally. |
| Extended Data Fig. 7 | **Cross-task similarity predictions at the level of individual regions and large-scale brain networks.** | ExtendedDataFigure7.png | (A) Cosine similarity between vectorized cross-task similarity values from observed (empirically-estimated) and from connectivity-predicted estimates (range 0.13–0.99; p_connperm_ < 0.05 in 355/360 regions). Regions with no values did not survive FC shuffling permutation test. (B) Similar comparison of the network-level cross-task similarity patterns. Grey dots indicate null distributions derived from FC shuffling. Network abbreviations from Cole-Anticevic Brain Network Partition: VIS1: primary visual network, VIS2: secondary (higher-order) visual network, SMN: somato-motor network, CON: cingulo-opercular network, DAN: dorsal attention network, LAN: language network, FPN: fronto-parietal network, AUD: auditory network, DMN: default-mode network, PMULTI: posterior multimodal network, VMM: ventral multimodal network, ORA: orbito-affective network. |
| Extended Data Fig. 8 | **Representational dimensionality is predictive of behavioral measures of cognitive flexibility.** | ExtendedDataFigure8.png | A) Average representational dimensionality across regions was positively correlated with cognitive flexibility scores across subjects. B) Similar relationship in individual regions (following multiple comparison corrections) was reflected in positive correlations in different regions. C) Dimensionality values averaged across sensory, association and motor regions were predictive of flexibility scores in a cross-validated (leave one subject out) manner. D) Coefficients from the cross-validation approach show higher values for sensory and association regions’ dimensionality values compared to motor regions in explaining the flexibility scores. E) The representational dimensionality values in the new dataset also follow compression-then-expansion pattern (average values across subjects). F) Relative compression in association regions compared to sensory and motor regions was reflected in almost all subjects, measured using compression ratio. These findings address an apparent paradox: although association regions show representational compression on average (Fig. 2), individual differences in absolute dimensionality within this organization—particularly in sensory and association regions—predict fluid intelligence, suggesting the compression-then-expansion pattern reflects canonical task organization while absolute dimensionality reflects individual capacity for flexible cognition. |
| Extended Data Table. 1 | Cross-task similarity comparison between tasks and networks. | ExtendedDataTable1.pdf | Cross-task similarity comparison between tasks and networks. |
| Extended Data Table. 2 | **Correlations between representational expansion and RSA template term coefficients, observed and predicted.** | ExtendedDataTable2.pdf | Anatomical specificity of the four RSA template terms: 9/10 highest sensory coefficients in sensory regions, 7/10 highest cognitive coefficients in association regions, 7/10 highest motor coefficients in motor regions (Fig. 5B). Conjunction-term coefficients correlated with representational expansion (ΔD_Rep_) more strongly than cognitive or motor terms alone, and comparably to the sensory term. |

***Do not insert additional rows - total number of Extended Data items must not exceed 10.***

1. **Supplementary Information:**
A. **PDF Files**

**Complete the Inventory below for all additional textual information and any additional Supplementary Figures, which should be supplied in one combined PDF file.**

- **Row 1:** A combined PDF containing any Supplementary Text, Discussion, Notes, Additional Supplementary Figures, Supplementary Protocols, simple tables, and all associated legends. **Only one such file is permitted**.
- **Row 2:** Nature Research’s Reporting Summary; if previously requested by the editor, please provide an updated Summary, fully completed, without any mark-ups or comments. **(Reporting Summaries are not required for all manuscripts.)**

**Note:** Please do not include a title page within your Supplementary Information file - a cover sheet that includes the title of your paper and a hyperlink to it will be automatically added while preparing your manuscript for publication

| Item | Present? | Filename  Whole original file name including extension. i.e.: Smith_SI.pdf. The extension must be .pdf | A brief, numerical description of file contents.  i.e.: *Supplementary Figures 1-4, Supplementary Discussion, and Supplementary Tables 1-4.* |
| --- | --- | --- | --- |
| Supplementary Information | Yes | Chakravarthula_SI.pdf | Supplementary Table 1 |
| Reporting Summary | Yes | Chakravarthula_RS.pdf |  |
| Peer Review Information | Choose an item. | *OFFICE USE ONLY* |  |

B. **Additional Supplementary Files**

**Complete the Inventory below for all additional Supplementary Files that cannot be submitted as part of the Combined PDF.**

- Do not list Supplementary Figures in this table (see section 2A)
- Where possible, include the title and description within the file itself
- Spreadsheet-based tables & data should be combined into a workbook with multiple tabs, not submitted as individual files.
- Compressed files are acceptable where necessary. ZIP files are preferred.
- Please note that the *ONLY* allowable types of additional Supplementary Files are:

| - Supplementary Tables | - Supplementary Audio | - Supplementary Videos | - Supplementary Software |
| --- | --- | --- | --- |
| - Supplementary Code | - Supplementary Data, for example: - Source Data for Supplementary Figures - Raw NMR Data, Cryo-EM Data - Computational Data, Crystallographic Data, etc. | | |

| Type | Number  Each type of file (Table, Video, etc.) should be numbered from 1 onwards. Multiple files of the same type should be listed in sequence, i.e.: Supplementary Video 1, Supplementary Video 2, etc. | Filename  Whole original file name including extension. i.e.: *Smith_ Supplementary_Video_1.mov* | Legend or Descriptive Caption  Describe the contents of the file |
| --- | --- | --- | --- |
| Choose an item. |  |  |  |
| Choose an item. |  |  |  |
| Choose an item. |  |  |  |
| Choose an item. |  |  |  |
| Choose an item. |  |  |  |
| Choose an item. |  |  |  |

***Add rows as needed to accommodate the number of files.***

**3. Source Data**

**Complete the Inventory below for all Source Data files.**

- Acceptable types of Source Data for Main Figures and Extended Data Figures or Tables are:
  - Statistical Source Data
    - Plain Text (ASCII, csv, TXT) or Excel formats only
    - Either one file for each relevant Figure, or a single file containing all source data, with clearly named tabs for each Figure/Extended Data Figure item
  - Full-length, unprocessed gels or blots
    - JPG, TIF, or PDF formats only
    - One file for each relevant Figure containing all supporting blots and/or gels, or a single file with clearly labeled gels or blots for each Figure/ED Figure item
- Source Data for Supplementary Figures is not allowed. Instead:
  - Include Unprocessed Gels or Blots for Supplementary Figures as additional Supplementary Figures to the main Supplementary Information PDF file.
  - Include Statistical Source Data for Supplementary Figures as ‘Supplementary Data’ files and list them in section 2B.
  - Please see [this example of Source Data](https://www.nature.com/articles/s41591-019-0505-4) in a publication.

| Parent Figure or Table | Filename  Whole original file name including extension. i.e.: *Smith_SourceData_Fig1.xls,* or *Smith_ Unmodified_Gels_Fig1.pdf* | Data description  i.e.: Unprocessed western Blots and/or gels, Statistical Source Data, etc. |
| --- | --- | --- |
| Source Data Fig. 2 | Figure2_source_file.xlsx | Statistical Source Data |
| Source Data Fig. 3 | Figure3_source_file.xlsx | Statistical Source Data |
| Source Data Fig. 4 | Figure4_source_file.xlsx | Statistical Source Data |
| Source Data Fig. 5 | Figure5_source_file.xlsx | Statistical Source Data |
| Source Data Extended Data Fig./Table 1 | ExtendedDataFigure1_source_file.xlsx | Statistical Source Data |
| Source Data Extended Data Fig./Table 2 | ExtendedDataFigure2_source_file.xlsx | Statistical Source Data |
| Source Data Extended Data Fig./Table 3 | ExtendedDataFigure3_source_file.xlsx | Statistical Source Data |
| Source Data Extended Data Fig./Table 4 | ExtendedDataFigure4_source_file.xlsx | Statistical Source Data |
| Source Data Extended Data Fig./Table 5 | ExtendedDataFigure5_source_file.xlsx | Statistical Source Data |
| Source Data Extended Data Fig./Table 6 | ExtendedDataFigure6_source_file.xlsx | Statistical Source Data |
| Source Data Extended Data Fig./Table 7 | ExtendedDataFigure7_source_file.xlsx | Statistical Source Data |
| Source Data Extended Data Fig./Table 8 | ExtendedDataFigure8_source_file.xlsx | Statistical Source Data |
