## Supplementary Data Table 1 for "Network geometry shapes the transformation of task representations across human cortex"

**Supplementary Information**

| Task | # Conditions | Task Condition | Key | stimType1 | stimType2 | stimType3 |
| --- | --- | --- | --- | --- | --- | --- |
| IAPS-Affective | 2 | Affective_pleasant | L_I | image | scene |  |
|  |  | Affective_unpleasant | L_M |  |  |  |
| IAPS-Emotional | 2 | Emotional_happy | R_I |  | face |  |
|  |  | Emotional_sad | R_M |  |  |  |
| Mental Rotation | 6 | mentalRotation_doesnotmatch_0_deg | R_M |  | objects |  |
|  |  | mentalRotation_doesnotmatch_50_deg | R_M |  |  |  |
|  |  | mentalRotation_doesnotmatch_150_deg | R_M |  |  |  |
|  |  | mentalRotation_match_0_deg | R_I |  |  |  |
|  |  | mentalRotation_match_50_deg | R_I |  |  |  |
|  |  | mentalRotation_match_150_deg | R_I |  |  |  |
| Object n-back | 12 | nBackPic_notpossible2backfork.jpg | none |  |  | fork image |
|  |  | nBackPic_possible2backfork.jpg | R_I |  |  |  |
|  |  | nBackPic_notpossible2backhydrant.jpg | none |  |  | hydrant image |
|  |  | nBackPic_possible2backhydrant.jpg | R_I |  |  |  |
|  |  | nBackPic_notpossible2backlamp.jpg | none |  |  | lamp image |
|  |  | nBackPic_possible2backlamp.jpg | R_I |  |  |  |
|  |  | nBackPic_notpossible2backplate.jpg | none |  |  | plate image |
|  |  | nBackPic_possible2backplate.jpg | R_I |  |  |  |
|  |  | nBackPic_notpossible2backwhistle.jpg | none |  |  | whistle image |
|  |  | nBackPic_possible2backwhistle.jpg | R_I |  |  |  |
|  |  | nBackPic_notpossible2backzip.jpg | none |  |  | zip image |
|  |  | nBackPic_possible2backzip.jpg | R_I |  |  |  |
| Concrete Permuted Rule Operations (CPRO) | 8 | CPRO_doesnotfollowRule_buttonOne | L_I | [image, text] | [object, word] | press button One |
|  |  | CPRO_followsRule_buttonOne | L_M |  |  |  |
|  |  | CPRO_doesnotfollowRule_buttonTwo | L_M |  |  | press button Two' |
|  |  | CPRO_followsRule_buttonTwo | L_I |  |  |  |
|  |  | CPRO_doesnotfollowRule_buttonThree | R_M |  |  | press button Three' |
|  |  | CPRO_followsRule_buttonThree | R_I |  |  |  |
|  |  | CPRO_doesnotfollowRule_buttonFour | R_I |  |  | press button Four' |
|  |  | CPRO_followsRule_buttonFour | R_M |  |  |  |
| Go / No-Go | 2 | GoNoGo_Go | L_I | text | word |  |
|  |  | GoNoGo_NoGo | none |  |  |  |
| Word Prediction | 3 | Prediction_random | L_M |  |  |  |
|  |  | Prediction_valid | L_I |  |  |  |
|  |  | Prediction_violation | L_M |  |  |  |
| Stroop | 8 | cong_color1 | L_M |  |  | ink color 1 |
|  |  | incong_color1 | L_M |  |  |  |
|  |  | cong_color2 | L_I |  |  | ink color 2 |
|  |  | incong_color2 | L_I |  |  |  |
|  |  | cong_color3 | R_I |  |  | ink color 3 |
|  |  | incong_color3 | R_I |  |  |  |
|  |  | cong_color4 | R_M |  |  | ink color 4 |
|  |  | incong_color4 | R_M |  |  |  |
| Theory of Mind | 4 | ToM_ToM_expected_false | L_M |  | sentence |  |
|  |  | ToM_ToM_expected_true | L_I |  |  |  |
|  |  | ToM_control_expected_false | L_M |  |  |  |
|  |  | ToM_control_expected_true | L_I |  |  |  |
| Arithmetic | 4 | DigitJudgement_target_absent | R_M |  | letter-digit |  |
|  |  | DigitJudgement_target_present | R_I |  |  |  |
|  |  | Math_equation_correct | R_I |  |  |  |
|  |  | Math_equation_incorrect | R_M |  |  |  |
| Motor Sequence | 5 | motorSequence_complex | all |  |  |  |
|  |  | motorSequence_simple_digit1 | L_M |  |  |  |
|  |  | motorSequence_simple_digit2 | L_I |  |  |  |
|  |  | motorSequence_simple_digit3 | R_I |  |  |  |
|  |  | motorSequence_simple_digit4 | R_M |  |  |  |
| Verbal n-back | 6 | nBack_notpossible2backA | none |  |  | letter 'A' |
|  |  | nBack_possible2backA | L_I |  |  |  |
|  |  | nBack_notpossible2backB | none |  |  | letter 'B' |
|  |  | nBack_possible2backB | L_I |  |  |  |
|  |  | nBack_notpossible2backC | none |  |  | letter 'C' |
|  |  | nBack_possible2backC | L_I |  |  |  |
| Response Alternatives | 12 | respAlt_1_prime_targPos1 | L_M | [text, array] | letter-digit |  |
|  |  | respAlt_1_prime_targPos2 | L_I |  |  |  |
|  |  | respAlt_1_prime_targPos3 | R_I |  |  |  |
|  |  | respAlt_1_prime_targPos4 | R_M |  |  |  |
|  |  | respAlt_2_primes_targPos1 | L_M |  |  |  |
|  |  | respAlt_2_primes_targPos2 | L_I |  |  |  |
|  |  | respAlt_2_primes_targPos3 | R_I |  |  |  |
|  |  | respAlt_2_primes_targPos4 | R_M |  |  |  |
|  |  | respAlt_4_primes_targPos1 | L_M |  |  |  |
|  |  | respAlt_4_primes_targPos2 | L_I |  |  |  |
|  |  | respAlt_4_primes_targPos3 | R_I |  |  |  |
|  |  | respAlt_4_primes_targPos4 | R_M |  |  |  |
| Spatial Map | 12 | spatialMap_1_element_targNum1 | L_M |  |  |  |
|  |  | spatialMap_1_element_targNum2 | L_I |  |  |  |
|  |  | spatialMap_1_element_targNum3 | R_I |  |  |  |
|  |  | spatialMap_1_element_targNum4 | R_M |  |  |  |
|  |  | spatialMap_4_elements_targNum1 | L_M |  |  |  |
|  |  | spatialMap_4_elements_targNum2 | L_I |  |  |  |
|  |  | spatialMap_4_elements_targNum3 | R_I |  |  |  |
|  |  | spatialMap_4_elements_targNum4 | R_M |  |  |  |
|  |  | spatialMap_7_elements_targNum1 | L_M |  |  |  |
|  |  | spatialMap_7_elements_targNum2 | L_I |  |  |  |
|  |  | spatialMap_7_elements_targNum3 | R_I |  |  |  |
|  |  | spatialMap_7_elements_targNum4 | R_M |  |  |  |
| Visual Search | 6 | visualSearch_target_absent_setsize12 | L_M |  |  |  |
|  |  | visualSearch_target_absent_setsize4 | L_M |  |  |  |
|  |  | visualSearch_target_absent_setsize8 | L_M |  |  |  |
|  |  | visualSearch_target_present_setsize12 | L_I |  |  |  |
|  |  | visualSearch_target_present_setsize4 | L_I |  |  |  |
|  |  | visualSearch_target_present_setsize8 | L_I |  |  |  |
| Biological Motion | 4 | BiologicalMotion_happy | R_I | video | point light |  |
|  |  | BiologicalMotion_sad | R_M |  |  |  |
|  |  | ScrambledMotion_fast | R_I |  |  |  |
|  |  | ScrambledMotion_slow | R_M |  |  |  |

### **Supplementary Data Table 1:** Task condition split of 16 active, visual tasks in Multi-domain Task Battery Dataset. For more information on task description, see the supplementary information of the original paper ^48^. Response key index: L_I: left index finger button, L_M: left middle finger button, R_I: right index finger button, R_M: right middle finger button, all: response using all four buttons was required, none: no response required.
