## Extended Data Table 2 for "Network geometry shapes the transformation of task representations across human cortex"

| Term | Observed r | Observed p | vs. conjunction (paired t) | Predicted r | Predicted p |
|---|---|---|---|---|---|
| Conjunction | 0.46 | 2.92e-07 | — | 0.34 | 0.006 |
| Sensory | 0.47 | 8.41e-10 | t(17)=-1.94, p=0.07 (n.s.) | 0.16 | 0.002 |
| Cognitive | 0.33 | 0.013 | t(17)=2.74, p=0.014 | 0.13 | 0.636 |
| Motor | -0.19 | 0.002 | t(17)=6.56, p=4.86e-06 | -0.15 | 0.039 |
