## Extended Data Table 1 for "Network geometry shapes the transformation of task representations across human cortex"

| Data Type | Task Comparison | Category | Mean VIS1 | Mean FPN | SD VIS1 | SD FPN | t Statistic | p-value | p (Bonferroni) | Cohen's d |
|---|---|---|---|---|---|---|---|---|---|---|
| Observed | Object N-back – Spatial Map | Shared Function (WM) | 0.2808 | 0.3347 | 0.0488 | 0.0476 | -10.4621 | $7.95 \times 10^{-9}$ | $4.77 \times 10^{-8}$ | -2.5374 |
| Observed | Verbal N-back – Spatial Map | Shared Function (WM) | 0.2212 | 0.3177 | 0.0530 | 0.0558 | -9.5790 | $2.90 \times 10^{-8}$ | $1.74 \times 10^{-7}$ | -2.3233 |
| Observed | Object N-back – Verbal N-back | Shared Function (WM) | 0.2773 | 0.3632 | 0.0618 | 0.0706 | -6.1485 | $1.07 \times 10^{-5}$ | $6.43 \times 10^{-5}$ | -1.4912 |
| Observed | Go No-Go – Arithmetic | Shared Stimulus (Word) | 0.4427 | 0.2807 | 0.0845 | 0.0498 | 11.9830 | $1.03 \times 10^{-9}$ | $6.17 \times 10^{-9}$ | 2.9063 |
| Observed | Go No-Go – Word Prediction | Shared Stimulus (Word) | 0.4819 | 0.2761 | 0.1036 | 0.0428 | 9.5515 | $3.02 \times 10^{-8}$ | $1.81 \times 10^{-7}$ | 2.3166 |
| Observed | Word Prediction – Arithmetic | Shared Stimulus (Word) | 0.4119 | 0.2482 | 0.0868 | 0.0438 | 9.4209 | $3.68 \times 10^{-8}$ | $2.21 \times 10^{-7}$ | 2.2849 |
| Model-Predicted | Verbal N-back – Spatial Map | Shared Function (WM) | 0.4612 | 0.5918 | 0.0780 | 0.0755 | -12.1001 | $8.86 \times 10^{-10}$ | $5.32 \times 10^{-9}$ | -2.9347 |
| Model-Predicted | Object N-back – Spatial Map | Shared Function (WM) | 0.5649 | 0.6363 | 0.0630 | 0.0580 | -5.7340 | $2.43 \times 10^{-5}$ | $1.46 \times 10^{-4}$ | -1.3907 |
| Model-Predicted | Object N-back – Verbal N-back | Shared Function (WM) | 0.5867 | 0.6634 | 0.0767 | 0.0719 | -4.4696 | $3.37 \times 10^{-4}$ | $2.02 \times 10^{-3}$ | -1.0840 |
| Model-Predicted | Go No-Go – Word Prediction | Shared Stimulus (Word) | 0.7982 | 0.4514 | 0.0766 | 0.0625 | 17.2661 | $3.26 \times 10^{-12}$ | $1.95 \times 10^{-11}$ | 4.1876 |
| Model-Predicted | Go No-Go – Arithmetic | Shared Stimulus (Word) | 0.7620 | 0.4592 | 0.0637 | 0.0489 | 16.1177 | $9.85 \times 10^{-12}$ | $5.91 \times 10^{-11}$ | 3.9091 |
| Model-Predicted | Word Prediction – Arithmetic | Shared Stimulus (Word) | 0.7323 | 0.4165 | 0.0889 | 0.0556 | 14.6785 | $4.36 \times 10^{-11}$ | $2.62 \times 10^{-10}$ | 3.5600 |
